# Unravelling the intraspecific variation in drought responses in seedlings of European black pine (*Pinus nigra* J.F. Arnold)

**DOI:** 10.1101/2025.10.20.683360

**Authors:** Muhammad Ahmad, Almuth Hammerbacher, Clara Priemer, Albert Ciceu, Marta Karolak, Sonja Mader, Sanna Olsson, Johann Schinnerl, Sebastian Seitner, Selina Schöndorfer, Paula Helfenbein, Jakub Jez, Michaela Breuer, Ana Espinosa-Ruiz, Teresa Caballero, Andrea Ganthaler, Stefan Mayr, Dominik K. Großkinsky, Stefanie Wienkoop, Silvio Schueler, Carlos Trujillo-Moya, Marcela van Loo

## Abstract

- Understanding intraspecific variation in drought tolerance is essential for predicting the adaptive capacity of forest species under climate change. Yet, the molecular basis of this variation remains poorly understood in ecologically and economically important conifers.
- We integrated high-throughput phenotyping with metabolomics and transcriptomics under standardized soil drying to investigate drought responses across nine climatically distinct provenances of the conifer *Pinus nigra*. We tested whether drought tolerance—measured as decline in maximum quantum yield of the photosystem II (Fv/Fm)—varies among provenances, follows a climatic cline, and involves trade-off with growth. To identify the underlying molecular basis, we performed metabolomics and transcriptomics in four provenances representing contrasting drought tolerance.
- Drought tolerance varied significantly among provenances and was decoupled from growth, yet showed no differentiation along the climatic cline. Drought tolerant provenances differed from sensitive ones in both constitutive and drought-induced levels of flavonoid and diterpene metabolites. Transcriptomic profiles further highlighted provenance-specific differences in gene expression related to flavonoids.
- Our results demonstrate the utility of integrating automated phenotyping with molecular profiling to uncover the metabolic basis of drought adaptation, laying the groundwork for targeted studies on metabolite function and tolerance strategies in non-model conifers.

## Introduction

Drought is an important environmental stressor limiting plant survival and productivity in both natural and managed ecosystems (Bartels & Sunkar, 2005; Fàbregas & Fernie, 2019). As climate change accelerates, drought events are becoming increasingly frequent, prolonged, and severe, posing a major threat to global ecosystem stability (Allen *et al*., 2015; Christian *et al*., 2021; Blackman *et al*., 2024). In conifers, extreme drought episodes have been linked to heightened forest mortality and declines in forest productivity (Wang *et al*., 2021; Hammond *et al*., 2022). This has prompted efforts to characterise inter- and intraspecific variation in drought tolerance of tree species (Isaac-Renton *et al*., 2018; Depardieu *et al*., 2020), define here as the ability to survive and grow under drought (Roskilly *et al*., 2025). The intraspecific variation constitutes an important reservoir of adaptive potential, with provenances adapted to drier or more variable climates offering potential sources of resilience. Understanding the extent of this variation, especially at early developmental stages, is critical for predicting adaptive potential, and informing strategies such as assisted migration, provenance selection, and conservation planning (Aitken & Whitlock, 2013; Franks *et al*., 2014; Aitken & Bemmels, 2016). Moreover, it can also reveal potential constraints or trade-offs—for instance, where drought tolerance may come at a cost to growth or other stress responses (Vanwallendael *et al*., 2019; Candido-Ribeiro & Aitken, 2024).

Provenance trials, in which growth responses are reconstructed from tree-ring measurements (Trujillo-Moya *et al*., 2018; Schueler *et al*., 2021), have long been the main approach to study genetic variation in drought tolerance. Although these trials remain important for understanding drought tolerance, they are constrained by high labor demands, uneven drought exposure, and confounding effects of environmental heterogeneity (Ahmad *et al*., 2025). By contrast, experimental drought can be applied more consistently under controlled conditions in seedlings, enabling the assessment of genetic variation in traits relevant to drought tolerance that are difficult to capture in adult trees (Roskilly *et al*., 2025). Among such traits, dark-adapted chlorophyll fluorescence (Fv/Fm, the maximum quantum yield of the photosystem II (PSII)) has emerged as particularly informative (Candido-Ribeiro & Aitken, 2024; Roskilly *et al*., 2025). Fv/Fm exhibits greater inter- and intraspecific variability than other traits such as cavitation resistance or rehydration capacity (Lamy *et al*., 2014; Trueba *et al*., 2019). By contrast, under non-stressed conditions, Fv/Fm values are remarkably consistent across plant lineages, with typical values around 0.83 (Murchie & Lawson, 2013). This stability facilitates direct comparisons among individuals, provenances, and species— unlike growth traits, which often differ in their baseline values. Although declines in Fv/Fm represent a relatively late stress signal (Hu *et al*., 2023), they provide an objective quantification of irreversible damage and have been identified as strong predictors of survival under drought in conifers and other species (Woo *et al*., 2008; Garcia-Forner *et al*., 2016; Guadagno *et al*., 2017). Furthermore, because it can be measured rapidly and at high throughput, Fv/Fm represents a practical and scalable trait for detecting genetic variation in drought tolerance.

While studies on intraspecific variation in drought tolerance are increasingly emerging in conifer seedlings (Bansal *et al*., 2015; Csilléry *et al*., 2020; Candido-Ribeiro & Aitken, 2024), molecular understanding of the mechanisms underlying this variation continues to lag behind in conifers. This gap is particularly pronounced in ecologically important, yet genetically less tractable, non-model conifers, where the limited molecular insight constrains the development and application of molecular markers for screening and selecting provenances for drought tolerance.

Most current knowledge of drought responses in contrasting drought tolerant genotypes or provenances at the molecular level stems from studies in angiosperms, particularly model species such as Arabidopsis and major crops (Turner, 2018; Zhang *et al*., 2024). Studies in these systems have shown that plants employ a combination of strategies across multiple levels to tolerate drought, a substantial fraction of which operate directly at the metabolic level (Schrieber *et al*., 2023). Together, these responses determine a plant’s ability to maintain physiological function and survive under water deficit. Key metabolites, revealed by targeted and/or untargeted metabolomics include proline and soluble sugars, which contribute to osmotic regulation; abscisic acid (ABA), a central signal in stomatal regulation and drought-induced transcriptional responses; and flavonoids and terpenoids, which serve antioxidant and membrane-stabilizing functions (Fàbregas & Fernie, 2019; Tiedge *et al*., 2022). Additionally, xanthophyll-cycle pigments (e.g., zeaxanthin) enhance photoprotection by dissipating excess excitation energy and mitigating photooxidative damage (Jahns & Holzwarth, 2012). While several of these compounds have been associated, albeit to varying extends, to drought tolerance in model plants (Szabados & Savouré, 2010; Vaughan *et al*., 2015; Turner, 2018; Fàbregas & Fernie, 2019; Tiedge *et al*., 2022; Zhang *et al*., 2023a), their contribution to variation in drought tolerance in non-model plants, especially in conifers, remains largely unexplored.

High-throughput plant phenotyping (HTPP) combined with transcriptomic and metabolomic profiling offers a powerful framework to characterize intraspecific variation in drought responses across multiple biological levels. When applied across diverse provenances, these approaches can uncover both the variation in drought tolerance and the molecular mechanisms that underpin it (Li *et al*., 2020; Lou *et al*., 2025). However, meaningful comparisons in drought stress experiments require that provenances experience identical levels of water deficit —a challenging task due to the strong relationship between water loss, plant size, and transpiration rate (Juenger & Verslues, 2023; Moshelion *et al*., 2024). This confounding effect can obscure true variation in drought tolerance and underlying mechanisms (Moshelion *et al*., 2024). The use of same-aged seedlings and recent advances in automated phenotyping platforms and gravimetric irrigation systems now allow precise soil moisture control (Paul *et al*., 2019; Langan *et al*., 2024) and robust, high-throughput assessment of drought responses across large cohort of provenances. Analyzing seedlings provides insights into a critical and sensitive life stage of trees, making the results directly relevant for natural regeneration, afforestation, and reforestation efforts under climate change.

Using a HTPP platform, we conducted a phenotyping experiment on European black pine (*Pinus nigra* J.F. Arnold), a widely distributed yet genetically fragmented conifer of ecological and economic importance across Europe (Vallauri *et al*., 2002; Thiel *et al*., 2012). Its broad geographic and climatic range (Caudullo *et al*., 2017) make *P. nigra* an ideal model for investigating intraspecific variation in drought tolerance in conifer trees. Although *P. nigra* is often considered for assisted migration in Central European forestry because of its high general drought tolerance, evidence of intraspecific variation in drought responses is inconsistent, with studies reporting both significant differences across provenances (Schirmer *et al*., 2022; Fkiri *et al*., 2024a,b) and lack of variation (Lebourgeois *et al*., 1998; Thiel *et al*., 2012). Moreover, little is known about the molecular responses and mechanisms underlying provenance-level differences in drought tolerance—particularly during the vulnerable seedling stage, when selection pressures are likely to be strongest (MacAllister et al., 2019).

To address these gaps, we applied HTPP under standardized drought conditions (Ahmad *et al.,* 2025) with a robotic gravimetric irrigation system to assess drought tolerance using Fv/Fm across seedlings from nine genetically diverse provenances. HTPP was followed by targeted and untargeted metabolomics and transcriptomics in provenances contrasting in drought tolerance. Specifically, we aimed to: (1) estimate interprovenance variation in drought tolerance and test its association with source climate; (2) identify conserved metabolic and transcriptional responses across provenances to reveal shared molecular signatures of drought adaptation in *P. nigra*, and (3) determine whether drought tolerance in contrasting provenances is linked to distinct patterns of metabolic and transcriptional profiles, thereby identifying putative molecular markers of drought tolerance.

## Materials and Methods

### Plant materials and conditions of growth

Seeds of nine provenances of *P. nigra* covering the geographic distribution of five subspecies according to Caudullo et al. (2017; see Fig. S1a; Table S1) were obtained from different sources. Since subspecies division is controversial (Olsson *et al*., 2020), we do not apply the subspecies concept in our manuscript. Seeds were sown in 250 mL pots (>3 seeds/pot), filled with 80 g of substrate (dry weight: 39 ± 1 g; Gramoflor Topf + TonXL). The pots were watered to ∼ 50% of soil volumetric water contents (SVWC) and were covered with transparent plastic domes to maintain 100% relative humidity. Plants were kept under these conditions for 14 days, and during this period, the temperature was 21°C, the light intensity was 100 μmol m^-2^ s^-1^ photosynthetic photon flux density (PPFD; at the substrate level), and light: dark was cycled for 16:8 h. After 14 days, when the seeds were germinated, domes were removed, seedlings were singularized to one per pot, and light intensity was gradually increased to 200 μmol m^-2^ s^-1^ PPFD. Relative humidity was set to 60%, while other environmental conditions were maintained as described above. All plants were kept in a phytotron of the Plant Science Facility (VBCF Vienna BioCenter Core Facilities GmbH, Vienna, Austria) until HTPP.

### Drought treatment and sampling

Six weeks after sowing, SVWC was gradually reduced to ∼13% (Fig. S1b), corresponding to a soil water potential (ψ_s_) of −0.25 MPa according to the water retention curve of the substrate (Ahmad *et al*., 2025). At this stage, each provenance consisted of 30 seedlings (except PN9 with 18), with half of the seedlings randomly assigned to a drought (D) and half to a well-watered (WW) treatment. For the D treatment, SVWC was further reduced to 7% (ψ_s_ = −3.0 MPa) over a 6-day period of progressive drought and then maintained at this level for additional 12 days. WW plants were watered to ∼ 30% SVWC (ψ_s_ = −0.02 MPa) and maintained at this level throughout the rest of the experiment (Fig. S1b). The drought severity here was comparable to a previous study of (Lebourgeois *et al*., 1998) on *P. nigra.* At day 18 (the final time point), shoots with needles were cut above the stem and flash-frozen in liquid nitrogen. For metabolite and mRNA-seq analyses, needles from 3-4 plants per treatment and provenance were randomly pooled to form one biological replicate, with 3-4 biological replicates per provenance and treatment were used in total.

### High-throughput plant phenotyping

Automated HTPP was initiated four days prior to the onset of the D treatment at the PHENOPlant (Plant Science Facility, VBCF Vienna BioCenter Core Facilities GmbH). The facility is equipped with a custom designed PSI PlantScreen™ Modular System (Photon System Instruments spol. s r.o., Drasov, Czech Republic). Over a period of 22 days (4 days prior the onset of D and 18 afterwards), seedlings of both groups were phenotyped on 16 days (time points). Plants were watered twice daily using an automated weighing and watering station to maintain the predefined SVWC levels.

Before phenotyping, manual support to the stem was provided by inserting blue sticks in the pot along the stem. Plants were loaded in multi-well trays (5 columns x 4 rows configuration). To avoid overlap between plants, each tray was only loaded with 6 pots randomly assigned to each of the six positions. A total of 43 trays containing 258 seedlings (n = 6/tray) were then randomly distributed on 2 lanes of the phenotyping centre. Trays were automatically loaded onto the imaging platform and imaged from the top (top-view) using chlorophyll fluorescence (Chlf), and RGB imaging sensors (Fig. S1c) to measure Fv/Fm and canopy area (projected green area in mm^2^ of the plant from top view), respectively.

Fv/Fm was measured on dark adapted plants (≥30 min) using a FluorCam FC-800MF camera (Photon System Instruments spol. s r.o., Drasov, Czech Republic). RGB images were captured using a PSI RGB camera (12.36-megapixel CMOS sensor, Sony MX253LQR-C) with a Samyang 16 mm f2 AS UMC CS lens. Image analysis and segmentation was performed as described in Supplementary materials (Methods S1).

### Statistical analysis of phenotypic data

We modelled Fv/Fm using a Generalized Additive Mixed Model (GAMM; (Wood, 2017)). Provenance was included as a parametric term, and replicate was treated as a random effect to account for autocorrelation. Time was modelled as a smooth term with 12 knots using thin-plate regression splines, and model fitting was done using Restricted Maximum Likelihood (REML; Wood, 2017). To assess provenance effects, Akaike Information Criterion (AIC) values were compared between models with and without provenance as a fixed factor. To test whether Fv/Fm is influenced by plant size—given that larger plants may experience greater drought stress due to higher transpiration (Moshelion *et al*., 2024)—absolute canopy area was included as a covariate in additional model comparisons. Finally, to rank provenances for drought tolerance, percentage loss of chlorophyll fluorescence Fv/Fm (PLCF) at the final drought time point relative to well-watered controls was calculated as:

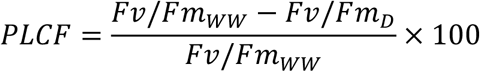

The PLCF metric reflects cumulative damage to PSII over the entire drought period. For simplicity, along the entire spectrum of PLCF, we refer to contrasting provenances with lower PLCF values (lower cumulative damage to PSII) as drought tolerant (DT) and those with higher values (greater cumulative damage to PSII) as drought sensitive (DS), irrespective of the underlying mechanisms.

For growth, we estimated canopy area increment (CAI) as the relative increase in canopy area compared to the first measurement (day −4) within each treatment and provenance, thereby reflecting growth relative to the initial size. In addition, relative canopy area increment (rCAI) was estimated to reflect the proportional growth reduction under drought compared to the well-watered plants for each provenance as:

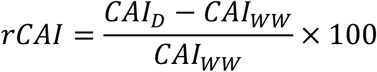

For PLCF and rCAI, one-way ANOVA was used to assess the effect of provenance, while two-way ANOVA was applied to CAI to test the effects of treatment, provenance, and their interaction at the final time point. Data were normalized using the bestNormalize R package to meet assumptions of normality.

To assess whether provenances with greater CAI under well-watered conditions were more drought tolerant, we regressed CAI against PLCF. A significant positive correlation would suggest a trade-off, where provenances investing more in canopy expansion under well-watered conditions exhibit lower drought tolerance.

To evaluate the relationship between climate of origin and drought tolerance as well as growth, we analysed the correlation between PLCF, CAI or rCAI and 19 bioclimatic variables retrieved from WorldClim (Hijmans *et al*., 2005). In addition, the 19 variables were summarised by principal component analysis (PCA), and the first two PCs were also included as predictors.

### Targeted metabolomics: assessment and analysis

We quantified 36 drought-related metabolites (Table S2-S3) known from conifers and other plant species (Savi *et al*., 2019; Zhang *et al*., 2024; Ahmad *et al*., 2025) using targeted analysis. Soluble carbohydrates, total phenolics, and condensed tannins were quantified using a sequential extraction protocol (Preiner *et al*., 2024), whereas proline was determined following (Carillo *et al*., 2008). Flavonoids (Table S3) were extracted and analysed following the method detailed in Supplementary materials (Methods S1, (Hammerbacher *et al*., 2018)). Xanthophylls (violaxanthin, neoxanthin, zeaxanthin, lutein), carotenoids (β-carotene -carotene, cis-β-carotene), chlorophylls (chl a, a’, b, b’) and tocopherols (α-, β- and γ-tocopherol) were extracted and quantified as describe in (Richins *et al*., 2014). Extraction and quantification of terpenes was performed using the method of (Joubert et al., 2023), and ABA using an untargeted metabolomics protocol (see the section below).

Metabolite data were log-transformed and analysed using two-tailed t-tests to compare treatments within each provenance, with p-values adjusted using the false discovery rate (FDR) method. Metabolites showing significant differences (FDR < 0.05) in at least one provenance were subsequently analysed by ANOVA, followed by Tukey’s HSD tests for pairwise comparisons among provenances when main effects were significant.

### Untargeted metabolomics: assessment and analysis

A Waters UHPLC coupled to a Waters SYNAPT G1 HDMS mass spectrometer was used to generate accurate mass data. For chromatographic separation a Waters HSS T3 C18 column (150 mm × 2.1 mm, 1.8 μm) was used at a temperature of 60°C and a binary solvent mixture consisting of water (A) containing 10 mM formic acid (pH 2.3) and acetonitrile (B) containing 10 mM formic acid. The initial conditions were 90% A at a flow rate of 0.4 mL min^−1^, maintained for 1 min, followed by a linear gradient to 1% A at 20 min and held for 2 min. One μL of the sample was injected. A SYNAPT G1 mass spectrometer was used in V-optics and operated in electrospray mode. Leucine enkephalin (50 pg mL^−1^) was used as reference calibrant to obtain mass accuracies between 1 and 5 mDalton (mDa). The mass spectrometer was operated in positive mode with a capillary voltage of 2.5 kV, the sampling cone at 30 V and the extraction cone at 4.0 V. The scan time was 0.1 s covering the 50 to 1200 Dalton mass range. The source temperature was 120°C and the desolvation temperature was set at 450°C. Nitrogen gas was used as the nebulization gas at a flow rate of 550 L h^−1^ and cone gas was added at 50 L h^−1^. MassLynx 4.1 (SCN 872) software was used for data acquisition.

The raw high-resolution mass spectrometer (HRMS) data was converted into .mzXML format using the peak-picking algorithm in MS convert in ProteoWizard (Adusumilli & Mallick, 2017) and uploaded to XCMS-online (Huan *et al*., 2017) to generate a feature table. The feature table was manually curated to include metabolites that eluted from the column between 2 and 16 minutes. In addition, only features, which were 50 times higher than the baseline, were included. The data was normalized by log transformation and range scaling. A PCA plot was generated. Based on PCA results, a feature table grouping provenances based on their tolerance was generated. The features were manually curated to remove isotopes, fragments and artefacts, and tentatively identified using Mass Bank (Horai *et al*., 2010), SIRIUS (Dührkop *et al*., 2019) and other resources (Vinaixa *et al*., 2016). A rarefied dataset with tentatively annotated features was used to create a heat map using the normalization settings above. An ABA standard curve from 10^-8^ to 10^-4^ mg/mL was prepared and analysed as above. Peaks with the same retention time and mass spectra were extracted from the sample HRMS chromatograms and quantified by using this standard curve. Statistical analysis was performed with two-way ANOVA to test the effects of provenance, treatment, and their interaction, followed by t-tests or Tukey’s HSD where main effects were significant.

### RNA-seq: RNA isolation, library preparation, sequencing, and data analysis

RNA isolation was performed as described in (Ahmad *et al*., 2025). Sequencing libraries were prepared using a QuantSeq 3’ mRNA-Seq Library Prep Kit REV for Illumina following the manufacturer’s instructions. Following library quality control, libraries were pooled in equimolar concentration and were sequenced using the Illumina NextSeq 2000 platform in SR100 mode at Lexogen GmbH (Austria, Vienna). The obtained reads were quality-controlled and trimmed for low-quality bases and sequencing adapters by employing cutadapt (v1.18; (Martin, 2011)). In the absence of the *P. nigra* genome, we mapped high-quality reads to phylogenetically related recently published *P. tabuliformis* genome (v1 (Niu *et al*., 2022)) using splice variant aligner STAR (v2.6.1a; (Dobin & Gingeras, 2015)). Finally, gene counts were performed using featureCounts (v1.6.4; (Liao *et al*., 2014)).

We next identified differentially expressed genes (DEGs) using DESeq2 (v1.18.1; (Love *et al*., 2014)). For each provenance, DEG were identified against the respective control samples (e.g., PN1_D vs PN1_WW). We considered genes as differentially expressed if |log₂FC| > 1 and FDR < 0.05. We then identified shared and unique DEGs. Gene ontology (GO) enrichment was performed using topGO and significance was assessed using Fisher’s exact test (v2.48.0; (Alexa & Rahnenfuhrer, 2023). GO terms were considered significantly enriched if P-values were lower than 0.01.

## Results

### Effect of drought on photosystem II efficiency

Over the course of drought, Fv/Fm in seedlings of each provenance remained close to the maximum (∼ 0.83) until day 3 (ψ_s =_ −1.3 MPa), after which values declined, reaching provenance-specific minima by approximately day 12 and remaining near these minima for the rest of the drought period (Fig. 1a). The GAMM model that included provenance as a fixed effect was strongly preferred (P < 0.0001) over the model without this factor (Table 1), indicating that the decline in Fv/Fm differed significantly among provenances. Including canopy area as a covariate did not improve model fit (Table 1), highlighting that absolute size differences did not influence Fv/Fm decline. Drought tolerance, estimated based on PLCF, varied significantly between provenances (P < 0.001). PN1 (Austria) and PN3 (Spain) exhibited lower PLCF values, suggesting comparatively greater drought tolerance, whereas PN6 (Croatia), PN4 (Cyprus), and PN7 (Austria) showed higher PLCF, indicating reduced tolerance (Fig. 1b).

**Fig. 1.**
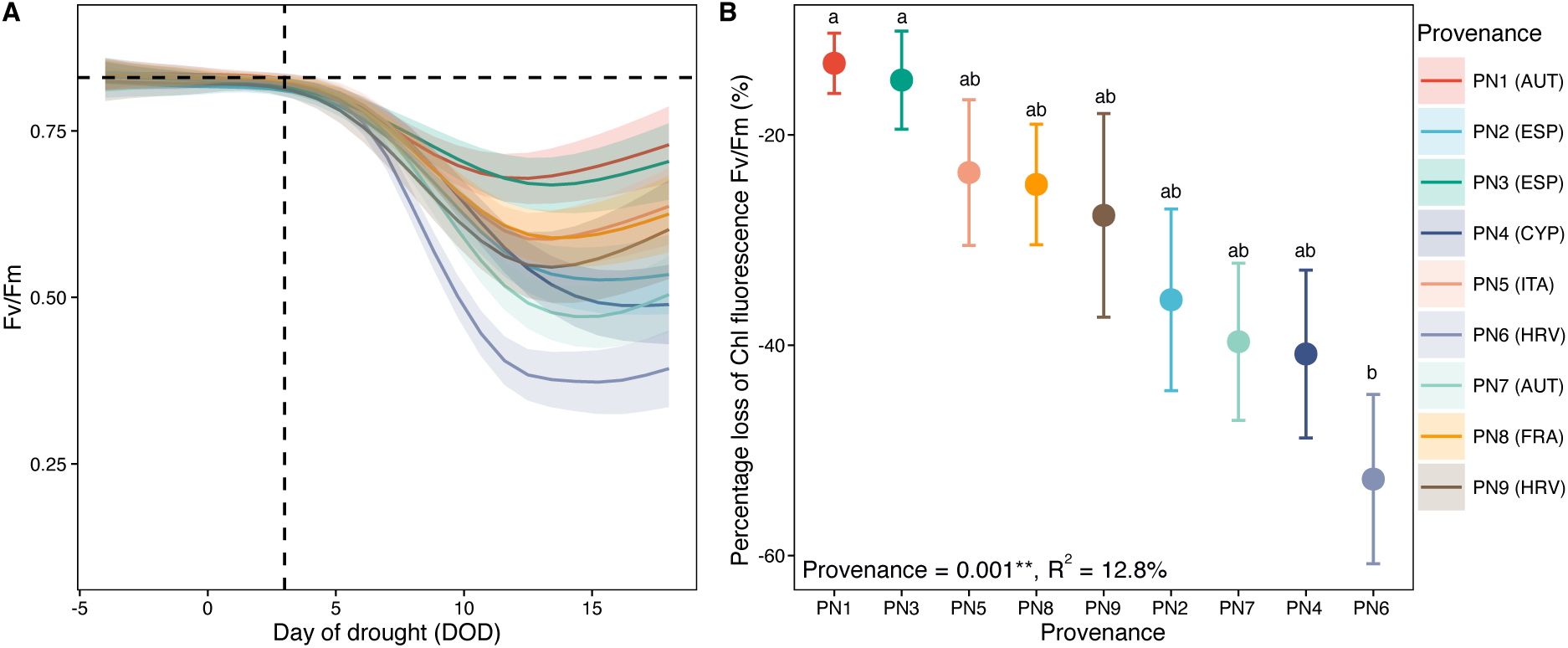
Drought effects on photosystem II (PSII) efficiency (Fv/Fm) and provenance differences in drought tolerance as assessed by percentage loss of chlorophyll fluorescence Fv/Fm (PLCF, %). (a) Trajectories of dark-adapted chlorophyll fluorescence (Fv/Fm) over the day of drought (DOD) for nine *Pinus nigra* provenances. Colored lines are GAMM fits; shaded bands are standard errors (n = 9 for PN9 and 14-15 for PN1-PN8). The vertical dashed line marks onset of decline in Fv/Fm (DOD = 3); the horizontal dashed line indicates the maximum Fv/Fm (∼0.83) under well-watered treatment average across all provenances. (b) Percentage loss of chlorophyll fluorescence Fv/Fm at the final drought time point (PLCF, %; relative to well-watered plants at DOD 18) for each provenance. Points show means ± SE (n = 9 for PN9 and 14-15 for PN1-PN8); letters denote Tukey groupings (α = 0.05). The provenance effect was significant (ANOVA: P < 0.001; R² = 12.8%). Lower PLCF values indicate that provenances maintained relatively higher PSII efficiency under drought (greater drought tolerance), whereas higher PLCF values reflect stronger declines in PSII efficiency (lower drought tolerance). Colors correspond to provenances. AUT: Austria, ESP: Spain, CYP: Cyprus, ITA: Italy, HRV: Croatia, FRA: France

**Table 1.**
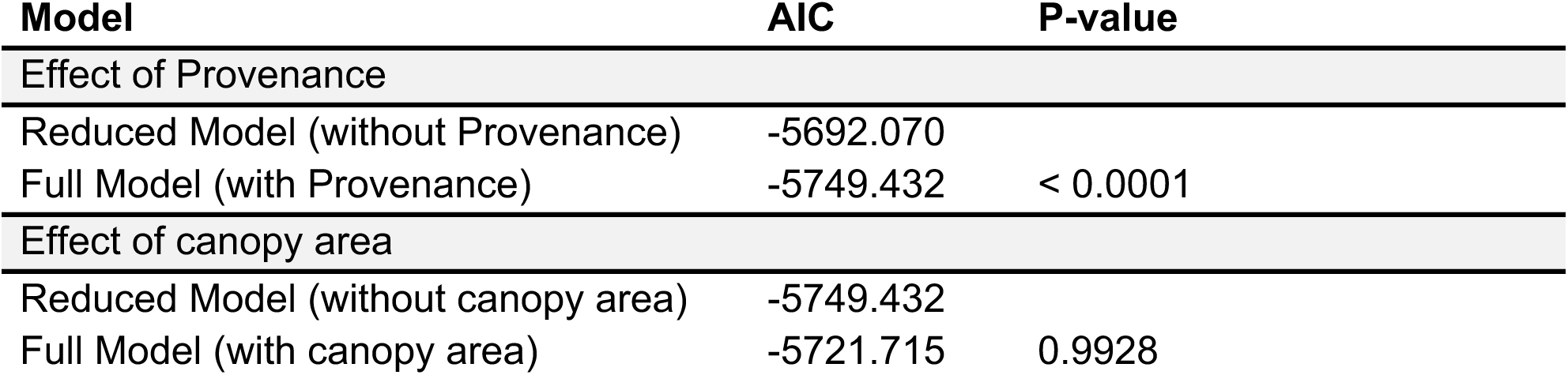
AIC comparison of models testing the effect of (i) provenance and (ii) canopy area on Fv/Fm.

### Effect of drought on growth

The drought treatment reduced growth in all provenances (Fig. 2). On average, CAI increased by ∼4% under D compared to ∼60% in the WW treatment (P < 2.2e^-16^), highlighting substantial effect of drought on growth. Moreover, provenances differed significantly in CAI (P < 4.4e^-07^), with PN1 exhibiting the highest CAI under both treatments (Fig. 2). The interaction between treatment and provenances was marginally significant (P = 0.075), suggesting that drought-induced reduction in CAI may vary among provenances. Provenance effects were likewise significant for rCAI, with PN6 (Croatia) showing the greatest relative reduction, whereas PN9 (Croatia) exhibited among the smallest declines (Fig. S2a).

**Fig. 2.**
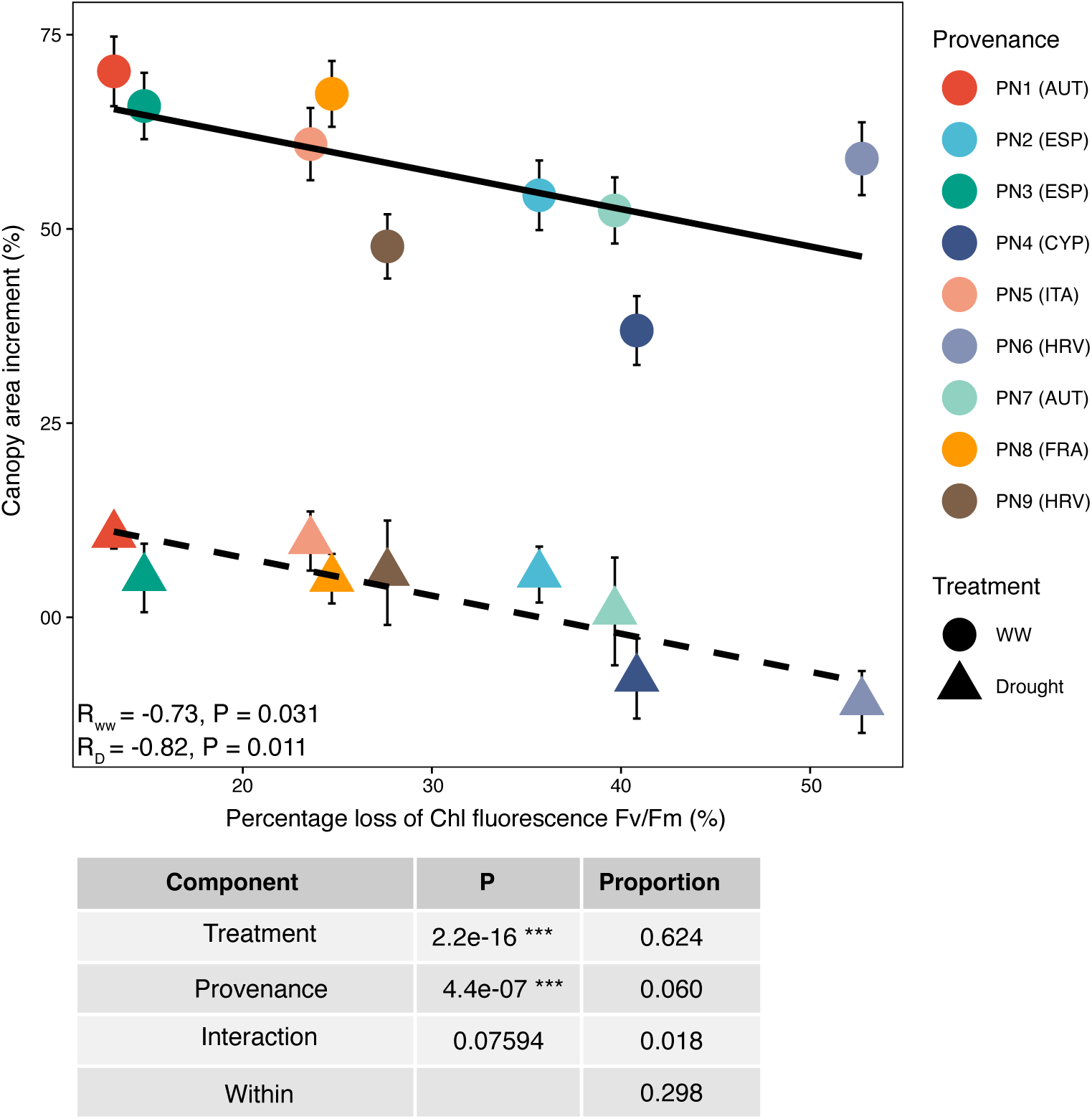
Effect of drought on growth and trade-off between growth and drought tolerance. Shown are canopy area increment (CAI, %) plotted against percentage loss of chlorophyll fluorescence Fv/Fm (PLCF, %) across nine provenances of *Pinus nigra* under well-watered (WW, circles) and drought (D, triangles) conditions. Each point represents provenance means ± SE (n = 9 for PN9 and 14-15 for PN1-PN8), with colors indicating provenances. Solid and dashed regression lines indicate relationships of CAI with PLCF under WW and D treatments, respectively, with corresponding correlation coefficients (R) and P-values shown. The variance partitioning table (two-way ANOVA) summarizes the proportion of variance in CAI explained by treatment (****), provenance (***), their interaction, and residuals. AUT: Austria, ESP: Spain, CYP: Cyprus, ITA: Italy, HRV: Croatia, FRA: France

### Lack of trade-off between growth and drought tolerance

There was a strong and significant negative relationship between PLCF and CAI under both WW (R = −0.73, P = 0.031) and D conditions (R = −0.81, P = 0.011; Fig. 2), suggesting that higher growth was associated with greater drought tolerance and thus no trade-off between these two traits. By contrast, although negative, the relationships between rCAI and CAI under well-watered conditions were non-significant (Fig. S2b), indicating that faster-growing provenances were not necessarily more vulnerable to drought.

### Clinal variation in drought tolerance of black pine along a climate gradient

To assess potential clines in drought tolerance across the climate gradient, we examined associations between the PLCF and bioclimatic variables. None of the correlations were significant. Similarly, climatic correlation with growth related variables (CAI or rCAI) were also weak and non-significant (Fig. S3), suggesting that no large-scale clinal variation of drought adaptation exists in black pine.

### Targeted metabolic profiling uncovers species-wide and provenance-specific drought responses

We next selected four extreme provenances—two drought sensitive (PN4, PN6; DS) and two droughts tolerant (PN1, PN3; DT)—to investigate the metabolic basis of contrasting drought performance. We hypothesized that metabolites increasing in response to D treatment represent common drought stress responses. In contrast, metabolites consistently higher in DT provenances, regardless of treatment may contribute to enhanced drought tolerance. To test this, we first employed a targeted approach and quantified metabolites known to be involved in drought responses, including ABA, proline, total soluble sugars, total phenolics, condensed tannins, carotenoids (n = 2), xanthophylls (n = 5), chlorophylls (n = 4), tocopherols (n = 3), terpenes (n = 3), and flavonoids (n = 14).

Among these compounds, 13 showed significant (FDR < 0.05) increases in at least one provenance in response to drought treatment (Fig. 3a,b, Table S4), with overall treatment effects also being significant across these metabolites (Table S5), suggesting species-wide drought-related responses. Moreover, provenances differed significantly (FDR < 0.05; Table S5) for these compounds except α-tocopherol and zeaxanthin; however, none exhibited consistently higher levels in DT compared to DS provenances (Fig. 3a). Instead, significantly higher levels were observed in the DS PN4 (proanthocyanidin B1) and PN6 (gallocatechin) or in both (kaempferol-3-O-glucoside-rhamnoside). Furthermore, PN3 and PN4 showed significantly higher concentrations of total phenolics, and condensed tannins compared to PN1 and PN6 (Fig. 3a).

**Fig. 3.**
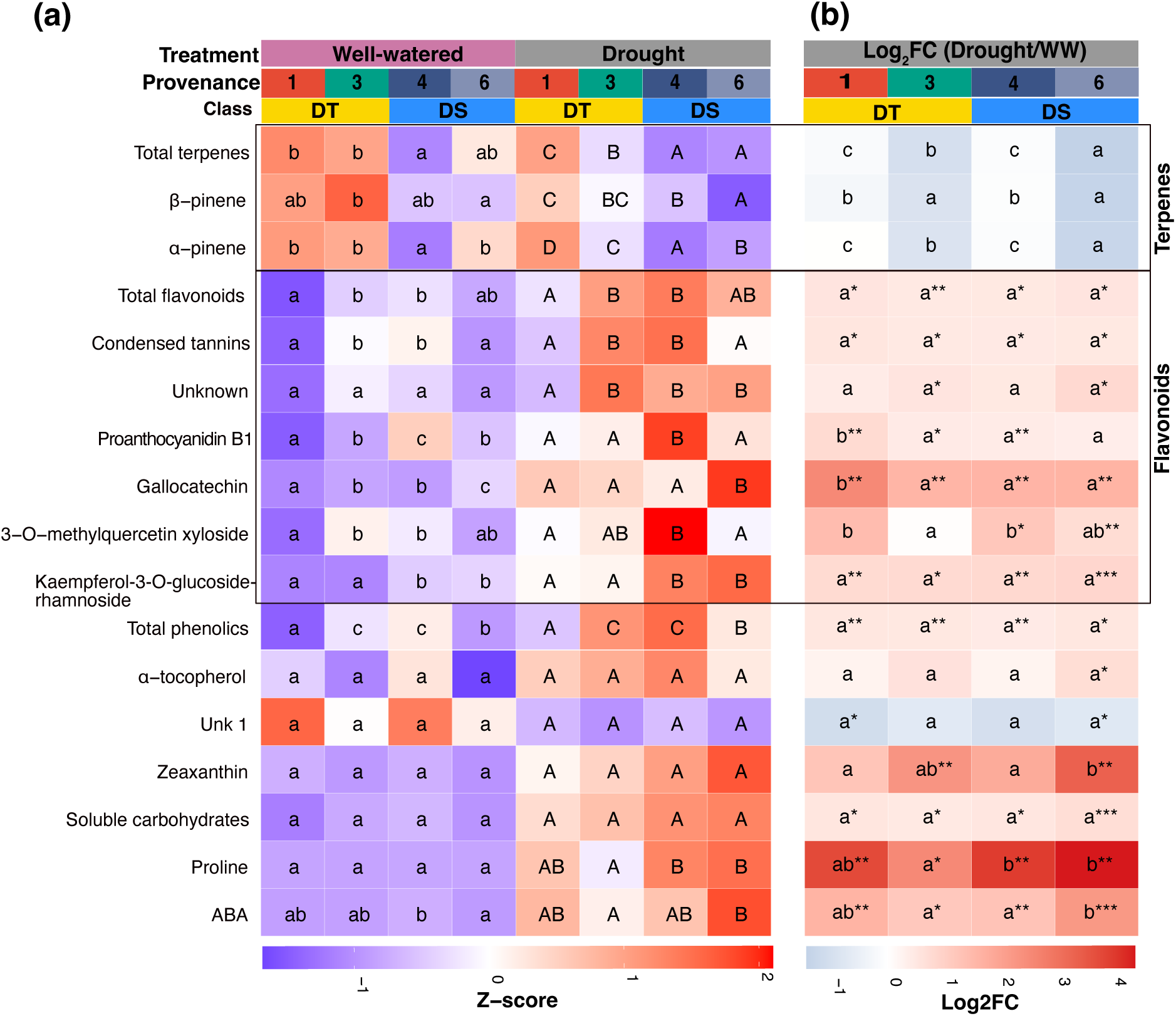
Shared and provenance-specific metabolic signatures of black pine (*Pinus nigra*) under well-watered (WW) and drought (D) conditions. (a) Heatmap of metabolites levels that showed significant changes (t-test; FDR < 0.05) in at least one provenance under D compared to WW treatment. Uppercase and lowercase letters indicate significant differences between provenance under D or WW conditions, respectively. Provenance-level differences within each treatment were assessed using ANOVA followed by Tukey’s HSD post hoc test (n = 3-4 biological replicates derived from the pool of 3–4 individuals per provenance). DT= Drought tolerant; DS = Drought sensitive. (b) Log₂fold changes (Log₂FC) of each metabolite, calculated as log₂(Drought / Well-watered). Asterisks indicate significant treatment effects within each provenance (t-test, FDR-corrected), while letters denote significant differences between provenances (Tukey’s HSD test). P-values ∗ = P < 0.05; ∗∗ = P < 0.01; ∗∗∗ = P < 0.001.

In terms of relative induction upon drought, most compounds were similarly upregulated across provenances (Fig. 3b). However, ABA, proline, and zeaxanthin exhibited significantly stronger fold-changes in the DS PN6 in at least one pairwise comparison. In contrast, the flavonoids gallocatechin and proanthocyanidin B1 showed markedly greater induction in the DT PN1 compared to all other provenances. These patterns suggest contrasting drought coping strategies: while PN3, PN4 and PN6 appear to rely on both constitutive levels and induced accumulation, PN1 primarily employ an inducible response for these metabolites.

Although no significant treatment effects were observed for terpenes—α-pinene, β-pinene, and total terpenes for any provenance, their overall absolute levels tended to be higher in DT provenances compared to DS ones under both conditions, with significant differences detected under drought (Fig. 3a).

### Untargeted profiling identifies putative metabolites linked to enhanced drought tolerance

To investigate additional metabolites involved in the drought stress response in black pine, we performed untargeted metabolomics, initially detecting 3205 metabolic features that were used in multivariate analysis. PCA revealed a complex picture with distinct baseline metabolic differences among provenances. Samples of each provenances clustered separately along PC1 (24.3% variance) and PC2 (20.7% variance), indicating that each provenance has a unique metabolic profile prior to D exposure (Fig. S4a). Since PC1 and PC2 were highly driven by constitutive differences between provenances, probably reflecting genetic differences, we focused on PC3 (13.7% variance) features which clustered provenances in the DT and DS groups under both treatments (Fig. S4b).

From PC3, 84 curated features were identified after the removal of fragments and isotopes. A new PCA and hierarchical clustering of these metabolic features revealed a clear separation between treatments along PC1, which explained 51.6% of the total variance (Fig. 4a). This separation was largely associated with the downregulation of metabolites in Cluster 1 in response to D, while a larger proportion of metabolites were induced and assigned to Cluster 2 (Fig. 4b). Nevertheless, the annotation of many of these metabolites remains unknown (Table S6).

**Fig. 4.**
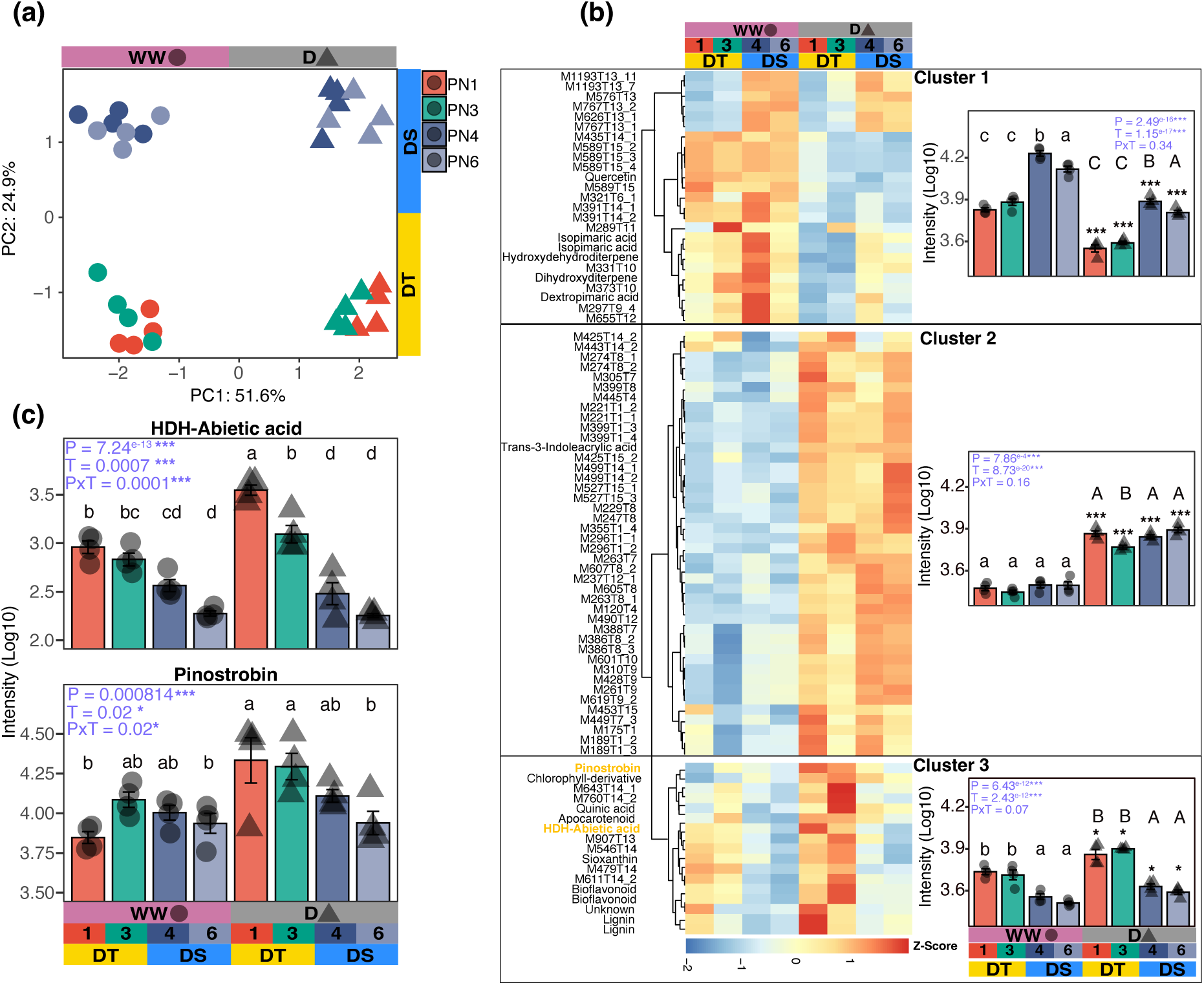
Metabolic signatures of drought tolerant (PN1 and PN3; DT) and drought sensitive (PN4 and PN6; DS) provenances of black pine (*Pinus nigra*) under well-watered (WW) and drought (D) treatment. (a) Principal component analysis of 84-curated compounds. (b) Heatmap and hierarchal clustering of 84 curated metabolites grouped into three distinct metabolic clusters. Inset barplots display average metabolite levels within each cluster. Error bars indicate the standard error of means (n = 4 biological replicates derived from the pool of 3–4 individuals per provenance and treatment). Asterisks above bars denote significant differences between treatments within a provenance (t-test; FDR-corrected). P-values ∗ = P < 0.05; ∗∗ = P < 0.01; ∗∗∗ = P < 0.001. Uppercase and lowercase letters indicate significant differences between provenances under D or WW conditions, respectively. Provenance-level differences were assessed using ANOVA followed by Tukey’s HSD post hoc test. Results of a two-way ANOVA are summarized in the upper right or left corner, indicating the effects of treatment (T), provenances (P), and their interaction (P × T) on average metabolite levels within each Cluster. Bold-yellow features indicate the putative hydroxydehydroabietic acid (HDH-Abietic acid) and pinostrobin discussed in the text. (c) Levels of HDH-Abietic acid and pinostrobin, both showing higher abundance in DT compared to DS provenances. Error bars indicate the standard error of means (n = 4 biological replicates derived from the pool of 3–4 individuals per provenance and treatment). Letters above bars indicate significant differences between provenances and treatment, determined by two-way ANOVA followed by Tukey’s HSD post hoc test. Results of a two-way ANOVA are summarized in the upper left corner, indicating the effects of treatment (T), provenance (P), and their interaction (P × T) on the respective metabolite.

Independent of treatment, provenances clustered distinctly along PC2 (24.9%), with the DT-group (PN1, PN3) separating from DS-group (PN4, PN6) (Fig. 4a). This pattern was consistent in both WW and D treatment, suggesting intrinsic metabolic differences existing between both groups and persisting under drought. These were primarily driven by Cluster 3 metabolites putatively annotated as quinic acid (phenolic precursor), pinostrobin (flavonoid), hydroxydehydroabietic acid (diterpenoid), apocarotenoid, adonoxanthin (tetraterpenoids), in addition to several unknown compounds. These metabolites were constitutively more abundant and relatively more induced under drought in DT provenances (Fig. 4b). Notably, hydroxydehydroabietic acid showed a significant provenance x treatment interaction (P < 0.0001), with higher basal levels in the DT group (log₂FC_DT/DS_ > 1), and under drought, its accumulation increased significantly more in DT (log₂FC_D/WW_ > 1) than in DS (log₂FC_D/WW_ = −0.13; 4B-C). Similarly, pinostrobin was significantly more induced in the DT provenance PN1, with PN3 reaching comparable levels under drought (Fig. 4b,c).

### Transcriptional profiling uncovers species-wide and provenance-specific drought responses

We next performed mRNA-seq to investigate the transcriptional basis of drought responses. PCA of gene expression separated WW and D treatments along PC1 (74.5% variance) and DT versus DS groups under drought along PC2 (7.5%), reflecting group-specific transcriptional differences (Fig. 5a). DT provenances exhibited approximately half the number of DEGs compared to the DS group (Fig. 5b; Data S1). Correspondingly, DEG counts were strongly negatively correlated with Fv/Fm (R² ≈ 1, P < 0.05; Fig. S5), linking transcriptional stability (less DEGs) to stable physiological performance under drought.

**Fig. 5.**
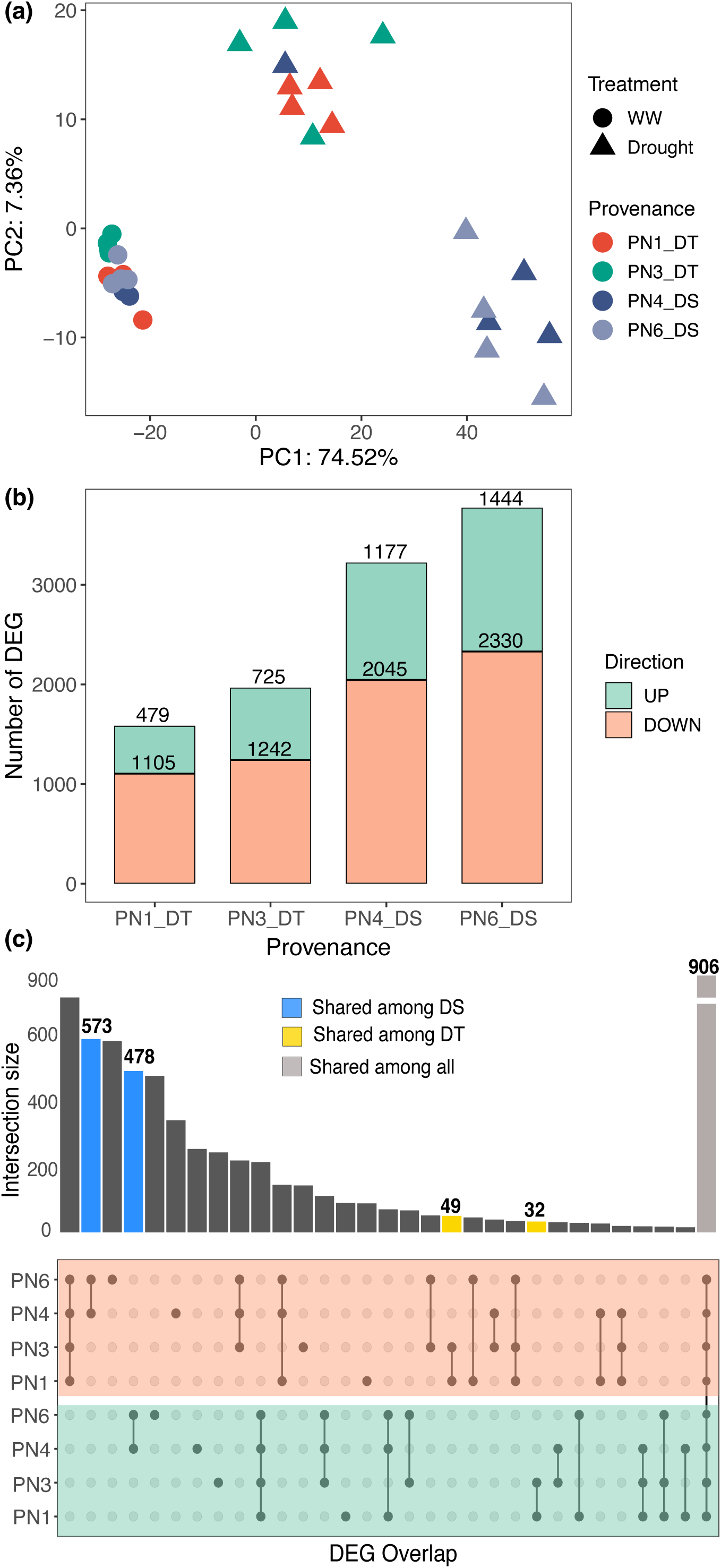
Transcriptional signatures of drought tolerant (PN1 and PN3; DT) and drought sensitive (PN4 and PN6; DS) provenances of black pine (*Pinus nigra*). (a) Principal component analysis (PCA) of expression profiles of four provenances of black pine under well-watered (WW) and drought treatment. (b) Number of differentially expressed genes (DEGs; |log2FC| > 1 and FDR < 0.05) in each provenance under drought relative to WW. (c) Overlap of DEG between provenances.

Both DT and DS provenances shared a conserved set of 906 drought-responsive genes (Fig. 5c; Table S7), representing a core component of the molecular drought-adaptation repertoire in *P. nigra*. This core set was enriched for GO terms related to osmotic stress, photoprotection, proanthocyanidin biosynthesis, photosynthesis, and water transport, among others (Fig. S6). Other notable annotations within this core set included genes involved in proline biosynthesis (P5CS), sugar transport (SWEET), ABA signalling (SNRK2, PYL, PP2C), and downstream ABA-responsive genes such as late embryogenesis abundant (LEA) proteins (Table S7).

We further assessed the overlap of DEGs within each group (Fig. 5c). DT provenances shared 5% of their total DEGs (32 upregulated, 49 downregulated; Data S1), suggesting a conserved transcriptional response underlying drought tolerance. Although several of these genes lacked functional annotations, the annotated subset included genes involved in histone modification (H3), calcium signalling (CIPK3), heat shock response (HSP20), and flavonoid biosynthesis (e.g., chalcone synthase [CHS] and anthocyanidin reductase [ANR]). In contrast, DS provenances shared ∼30% of their DEGs (478 upregulated, 573 downregulated; P < 0.05; Fig. 5c; Data S1). Upregulated shared genes in DS were enriched for sugar transport, phenylpropanoid metabolism, and oxidative stress responses (Table S8), whereas downregulated shared genes were, among others, linked to the photosynthetic machinery including PSII (e.g., psbS), PSI subunits (e.g., psaK), electron transport chain (e.g., petF), and ATP synthase genes-while these remained relatively stable in DT provenances (Fig. 6a).

**Fig. 6.**
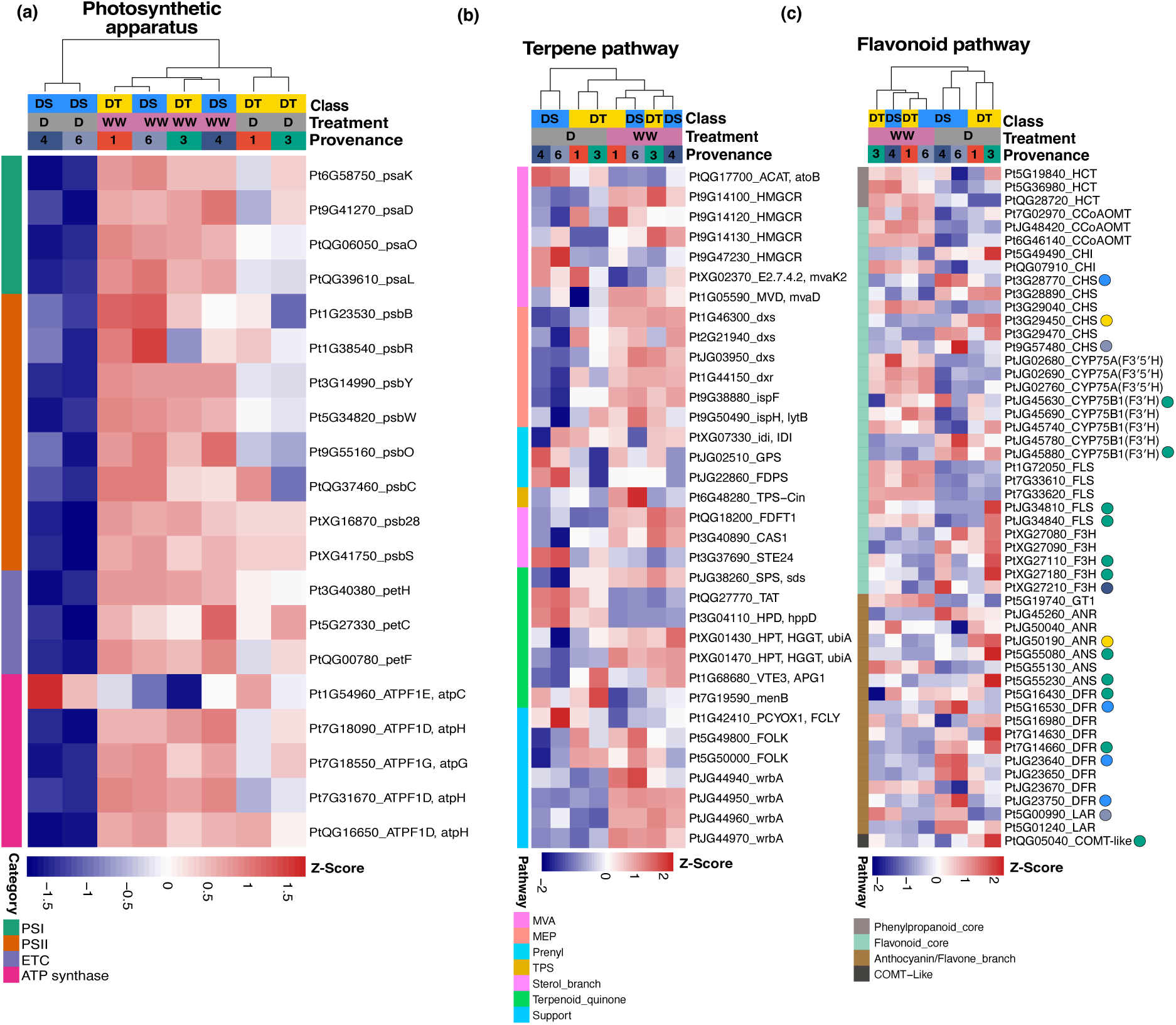
Transcriptional signatures of drought tolerant (PN1 and PN3, DT) and drought sensitive (PN4 and PN6, DS) provenances of black pine (*Pinus nigra*) across selected pathways. (a) Photosynthetic apparatus. Heatmap of z-score–scaled normalized expression values of genes encoding components of Photosystem I (PSI), Photosystem II (PSII), the electron transport chain (ETC), and the ATP synthase complex. Genes are grouped by functional category (row annotation). (b) Terpene biosynthesis pathway. Heatmap of z-score–scaled normalized expression values of genes involved in terpene biosynthesis. Only genes with FDR < 0.05 in at least one treatment comparison are shown. Rows are grouped into functional modules: mevalonate (MVA), methylerythritol phosphate (MEP), prenyltransferases, terpene synthase (TPS), pathway modifiers (sterol and terpenoid quinone) and pathway support. (c) Flavonoid biosynthesis pathway. Gene copies which are specifically expressed (log2FC >1 and FDR < 0.05) only in DT or in DS provenances or uniquely in each provenance are marked with color circles, respectively. Color scale for all panels indicates relative expression, from low (blue) to high (red).

### Transcriptomic regulation of terpene and flavonoid biosynthesis under drought in black pine

Given the observed differences in terpene- and flavonoid-related metabolites between treatments and provenance (Fig. 3 and 4c), we analysed the expression of potentially associated biosynthetic genes with significant differences (FDR < 0.05) in at least one provenance. Remarkably, based on expression levels of each of the pathway genes, DT and DS provenances grouped separately under D treatment (Fig. 6b,c), suggesting that modulation of terpene and flavonoid biosynthesis mirror broader transcriptional differences linked to drought performance. In the terpene pathway, DS provenances exhibited relatively stronger repression of genes from upstream MEP pathway, while DT provenances maintained relatively stable expression (Fig. 6b; Table S9). In contrast, flavonoid biosynthetic genes were modulated in both directions across all provenances, with distinct differences between DT and DS groups. For example, gene copies encoding CHS, flavanone hydroxylases (F3H, F3′H), flavonol synthase (FLS), dihydroflavonol 4-reductase (DFR), anthocyanidin synthase (ANS), anthocyanidin reductase (ANR), and caffeic acid-O-methyltransferase (COMT) being DEG differed between sensitivity groups or provenances (Fig. 6c; Table S10). These patterns suggest that higher drought tolerance might be associated with the preservation of terpene metabolism and a more targeted activation of flavonoid biosynthesis through distinct gene activation.

## Discussion

### Intraspecific variation in seedling drought responses: functional and molecular evidence

Empirical evidence for intraspecific variation in drought responses in *P. nigra* has been inconsistent. Earlier studies based on growth, survival and physiological traits reported no significant variation in drought responses at the seedling stage (Lebourgeois *et al*., 1998; Thiel *et al*., 2012), whereas more recent work has identified significant differences among provenances in both juvenile and adult trees (Schirmer *et al*., 2022; Fkiri *et al*., 2024a,b). Our findings align with the recent studies and extend them by showing that such variation is already detectable at the seedling stage. We observed a pronounced intraspecific variation in drought responses across *P. nigra* provenances, manifested at multiple biological levels—from growth and photosynthetic efficiency to transcriptomic and metabolic responses. Likewise, drought tolerance assessed based on PLCF, representing cumulative damage to PSII, showed significant variation between provenances.

Consistent with these findings, DS provenances showed strong downregulation of genes encoding the PSI/PSII core components, and key elements of the electron transport chain, while expression of these genes remained stable in the DT provenances, suggesting sustained carbon assimilation under water limitation in the latter. These molecular changes were associated with broader transcriptional and metabolic reprogramming: DS provenances exhibited nearly double the number of DEGs and more pronounced modulation of transcriptional profiles compared to DT provenances. These patterns are consistent with findings in other species—including switchgrass (Tiedge *et al*., 2022), sesame (You *et al*., 2019), wheat (Guo *et al*., 2025), African acacias (Weinheimer *et al*., 2025) and Norway spruce (Ahmad *et al*., 2025)— indicating increased sink activity for maintaining homeostasis under drought in DS. As all seedlings were grown under identical conditions, including substrate and soil moisture, the observed differences between provenances likely reflect underlying genetic variation. The emergence of these contrasts at the seedling stage suggests that provenance-specific drought responses are expressed early, potentially influencing seedling survival and, ultimately, demographic filtering in natural stands. Previous studies reporting no intraspecific variation at the seedling stage (Lebourgeois *et al*., 1998) (Thiel *et al*., 2012) likely underestimated the full range of variation within the species by focusing on a limited number of provenances from the central part of the species’ distribution. Moreover, environmental heterogeneity in field assessments (Lebourgeois *et al*., 1998; Thiel *et al*., 2012) may have further masked the differences. In contrast, our study combined comprehensive provenance sampling with rigorously controlled experimental conditions and multi-layered trait analyses, enabling robust differentiation of drought responses across provenances.

### No large-scale clinal adaptation to drought in *P. nigra*

While we found evidence for intraspecific variation in drought tolerance, associations between climate of origin and drought tolerance (as well as correlation with growth) were weak and non-significant, suggesting that no clinal variation in drought response along the geographic distribution of the species exists. Although we cannot rule out the possibility that stronger and more significant associations might emerge with broader and denser provenance sampling, our results are in line with previous studies on *P. nigra* (Tíscar *et al*., 2018; Santini *et al*., 2019), *P. pinaster* and *P. sylvestris* (Corcuera *et al*., 2011; Lamy *et al*., 2014) but differ from other pines (reviewed in (Ramírez-Valiente *et al*., 2022)). The lack of correlations might be explained by several factors. First, soil characteristics, which can buffer atmospheric drought stress (Cartwright *et al*., 2020), were not included as a predictor and may play an important role. Second, the fragmented distribution of *P. nigra* may have promoted genetic drift and reduced local adaptation signals, weakening climate–trait associations. Third, plasticity in physiological traits could enable provenances to cope with drought across a broad range of environments, obscuring adaptive signatures under controlled conditions. Fourth, conifers including *P. nigra* have experienced human translocations and planting outside their natural range, which may further blur patterns of local adaptation (Jansen *et al*., 2011; Vacek *et al*., 2023; Kovacs *et al*., 2024). Future studies that integrate soil characteristics and genetic analyses could shed more light on the drivers of drought tolerance and the role of local adaptation.

Collectively, our findings, and those reported by others, highlight that while the intraspecific variation in drought tolerance is evident, climate of origin is not a robust predictor for drought tolerance whilst selecting optimal provenances for assisted migration programs. Consequently, provenance response under drought should be assessed empirically rather than inferred solely based on climatic data. Simultaneously, the considerable phenotypic variation observed within and between provenances (Fig. 1,2) in our study suggests that significant genetic variation exists for selection of drought-tolerant breeding material for more resilient trees adapted to future climatic conditions. Importantly, the absence of a trade-off between growth and drought tolerance further indicates that selecting for higher tolerance may not necessarily compromise growth. This pattern has been previously observed in *P. nigra* and other conifer species (Ramírez-Valiente *et al*., 2022; Schirmer *et al*., 2022; Candido-Ribeiro & Aitken, 2024). However, tolerant provenances may exhibit proportionally larger reductions in growth, reflecting a strategy to minimize evapotranspiration and carbon demand under limited water supply.

### Shared and unique molecular signatures of drought responses in black pine provenances of contrasting drought tolerance

In contrast to angiosperms, the molecular basis of intraspecific variation in drought tolerance remains understudied in conifers, with relatively few examples reported in the literature (Nguyen-Queyrens & Bouchet-Lannat, 2003; Du *et al*., 2016; Kleiber *et al*., 2017a,b; Junker-Frohn *et al*., 2019). Through an in-depth investigation of *P. nigra* provenances with contrasting drought performance, we identified both shared and provenance-specific metabolic and transcriptional responses. Two major metabolite groups emerged: one broadly upregulated under drought across all provenances, indicating a general drought response across the species; the other consistently more abundant in DT provenances, suggesting a potential role in enhanced drought tolerance.

Among the first group of metabolites were ABA, proline, soluble carbohydrates, the xanthophyll zeaxanthin, total phenolics, flavonoids (kempferol-3-glucoside-rhamnoside, gallocatechin, total flavonoids, proanthocyandin B1), and several unidentified metabolites of the untargeted approach (Fig. 3,4). Their accumulation, together with the induction of genes in related biosynthetic pathways, points to a coordinated protective response involving osmotic adjustment, antioxidative defence, and hormonal regulation. For example, proline and soluble sugars contribute to osmotic adjustment—one of the main physiological mechanisms of drought adaptation in plants—by stabilizing proteins, preserving turgor, and mitigating dehydration-induced damage (Turner, 2018). Increased zeaxanthin, phenolics and flavonoids support photoprotection and antioxidant defence by dissipating excess light energy and scavenging reactive oxygen species, (Nakabayashi *et al*., 2014; Agati *et al*., 2020; Ferreyra *et al*., 2021; Changan *et al*., 2023) while elevated ABA levels reflect its central role in stomatal regulation (Gupta *et al*., 2020). Additional mechanisms by which ABA might support an effective protection from drought include the induction of protective proteins such as LEA proteins (Huang *et al*., 2018), which stabilize protein structures during dehydration (Goyal *et al*., 2005). Consistent with this, several LEA-encoding genes were upregulated in our transcriptomic data (Table S11). Activation of these responses across all provenances, together with the observed stabilization of Fv/Fm after 12 days, suggests that these metabolites helped to prevent further declines in photochemical efficiency. This supports the interpretation that they form a core component of the drought adaptation in *P. nigra*, enabling seedlings to maintain physiological function under sustained water deficit.

Among the second group of compounds were those assigned to cluster 3 in the untargeted metabolomic dataset (Fig. 4). Although the functional annotations and the mechanisms of actions of these compounds require further investigations, their higher abundance in DT provenances may contribute to the differences in drought performance of tested provenances. Their stronger accumulation under drought in DT, compared to DS provenances, support their role in conferring higher drought tolerance rather than involvement in general stress related responses. Among these compounds, we identified the methylated flavanone pinostrobin and the diterpene hydroxydehydroabietic acid as promising candidates which showed significant interactions between provenance and treatment.

Pinostrobin (Metsämuuronen & Sirén, 2019), is induced in response to beetle and fungal infection in different pine species (Fortier, 2022). It exhibits antioxidant properties (Patel *et al*., 2016), potentially supporting cellular protection during stress. Pinostrobin is synthesized from the flavanone pinocembrin via methylation, catalysed by O-methyltransferase enzymes (Chandran *et al*., 2022; Hanko *et al*., 2024). In our study, the differential expression of an O-methyltransferase related (COMT) gene between DS and DT groups, along with other flavonoid-related transcripts (Fig. 6c) may explain the observed differences in pinostrobin accumulation.

The second metabolite, potentially contributing to distinct drought performance, the hydroxydehydroabietic acid is an abietic acid derivative found in *P. nigra* and other pine species (Koutsaviti *et al*., 2017). It belongs to a class of diterpene resin acids that are abundant in conifers and serve defensive roles against pathogens and herbivores (Ro & Bohlmann, 2006). Drought-related roles of abietic acid derivatives and other diterpenes have been observed by several investigations in pine and other plant species. For example, the levels of dehydroabietic acid have been shown to increase in response to moderate drought stress in *P. sylvestris* (Turtola *et al*., 2003; Sancho-Knapik *et al*., 2017) and *P. elliottii* seedlings (Zhang *et al*., 2023b). Similarly, in crop plants (e.g., maize), higher levels of the diterpene kauralexin are associated to biotic and abiotic stress tolerance and mutants lacking in its synthesis were more sensitive to drought than the wildtype (Vaughan *et al*., 2015). More recently, elevated diterpene levels were correlated with higher drought tolerance of contrasting switchgrass genotypes (Tiedge *et al*., 2022). Tiedge et al. (2022) demonstrated a direct correlation between diterpene accumulation and transcript levels of pathway genes. In contrast, no such direct gene-to-metabolite relationships were observed in our study. Genes encoding abietadiene synthases—key enzymes in the biosynthesis of abietane-type diterpenes (Ro & Bohlmann, 2006)—were not differentially expressed under drought or between different groups. However, genes in the upstream MEP pathway maintained stable expression in DT provenances under drought, whereas they were strongly downregulated in DS provenances. Despite this transcriptional pattern, DT provenances accumulated significantly higher levels of hydroxydehydroabietic acid compared to DS. A similar pattern was observed for monoterpenes, which are also derived from the MEP pathway, where significantly higher metabolite levels were detected in DT provenances (Fig. 3). Together, these findings suggest that transcript abundance at the sampled time point may not fully capture final metabolite accumulation, which could instead reflect contributions from pre-existing metabolite pools, alternative biosynthetic routes, or regulatory events occurring at earlier growth or stress stages.

### Conclusions

Our study provides clear evidence of intraspecific variation in drought responses in *P. nigra*, detectable already at the seedling stage and manifested across functional, metabolic, and transcriptional levels. DS provenances exhibited stronger downregulation of growth and photochemical efficiency, accompanied by pronounced metabolic and transcriptional reprogramming when compared to the DT provenances—likely reflecting attempts to maintain physiological function. Integrated transcriptomic and metabolomic analyses revealed both shared core responses contributing to general drought adaptation and provenance-specific molecular signatures potentially underpinning enhanced tolerance. Beyond offering insights into intraspecific variation in drought responses and addressing links to growth and climatic origin, our findings identify a kaempferol derivative, pinostrobin, and hydroxydehydroabietic acid as potential metabolic markers, providing a foundation for future studies to investigate their role in drought tolerance of *P. nigra* and other conifers.

## Acknowledgements

We thank Florence Lee for her assistance in setting up and harvesting the experiment, Lenka Polonyova and Richarda Schuller for seed germination assay, Marlene Murauer, Charalambos Neophytou, Sotiris Sotiriou, Andreas Christou, Maurizio Sabatti, Tenente Colonnello, Silvia Biondini and Sanja Peric for providing seeds of black pine. We thank Ethan Stewart for performing image segmentation and analysis. We acknowledge The Austrian Research Promotion Agency (FFG) - R&D Infrastructure Funding Programme - PHENOPlant project #870446″ for the PHENOPlant research infrastructure. The Vienna BioCenter Core Facilities (VBCF) Plant Sciences Facility acknowledges funding from the Austrian Federal Ministry of Education, Science & Research; and the City of Vienna. We gratefully acknowledge the Dirección General de Biodiversidad, Bosques y Desertificación (Spain), under the Ministerio para la Transición Ecológica y el Reto Demográfico (MITECO), and particular the Centro Nacional de Recursos Genéticos Forestales El Serranillo (Spain), for providing the Spanish seed sources used in this study.

## Author contributions

MA, CTM and MvL conceived the study and designed the research together with SO. MA carried out the experiment with the support of SM, MB and MK and coordinated data collection. SSe, SSc, PH, and JJ conducted phenotyping. SMa and MK assisted with experimental work and sample preparation for metabolomics and RNA-seq. AH performed flavonoid, terpene, and untargeted metabolomics analyses. AER and TC conducted pigment and tocopherol measurements. DKG carried out proline analysis. CP, JS, and SW performed spectrophotometric measurements. AC conducted the GAMM analysis. SO contributed to seed source selection, study design, and performed functional gene annotation. SiSc, AG and StMa provided scientific and editorial advice. MA analysed the data and prepared the initial manuscript draft. CTM and MvL supervised the research and both shared the senior authorship equally. All authors contributed to manuscript editing, revision and approved the final version.

## Data Availability

Raw sequence data have been submitted to the NCBI Short Read Archive (SRA) under PRJNA1345943 and will be release upon publication.

https://dataview.ncbi.nlm.nih.gov/object/PRJNA1345943?reviewer=b4bugcd77k67iv1kocp6ipmb5u

---

**Supplementary figures and tables**

**Fig. S1-S6**

**Tables S1-S11** (provided as a separate excel file)

**Data S1** (provided as a separate excel file)

**Fig. S1.**
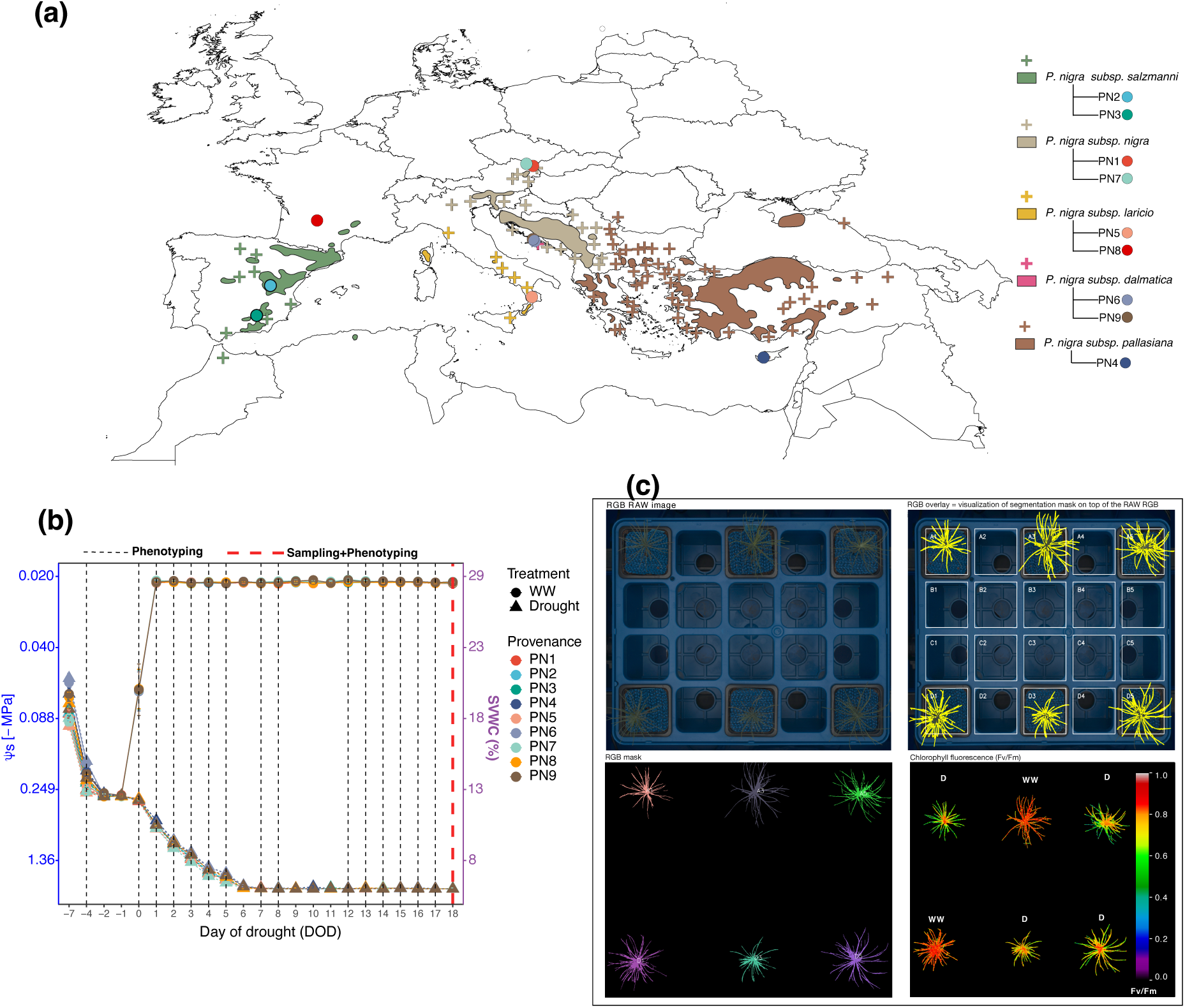
Study design. (a) Geographic origin of nine provenances representing five subspecies of black pine (*Pinus nigra*), shown alongside the natural distribution of each subspecies. (b) Drought progression measured as soil water potential over the 18-day drought stress experiment. Soil water potential decreased from −0.24 MPa to −2.97 MPa over six days and was maintained at this level for the remaining 12 days. (c) Schematic of the experimental set-up showing the imaging sensors used for high-throughput phenotyping. RGB: Red–Green–Blue imaging; Chlorophyll fluorescence imaging.

**Fig. S2.**
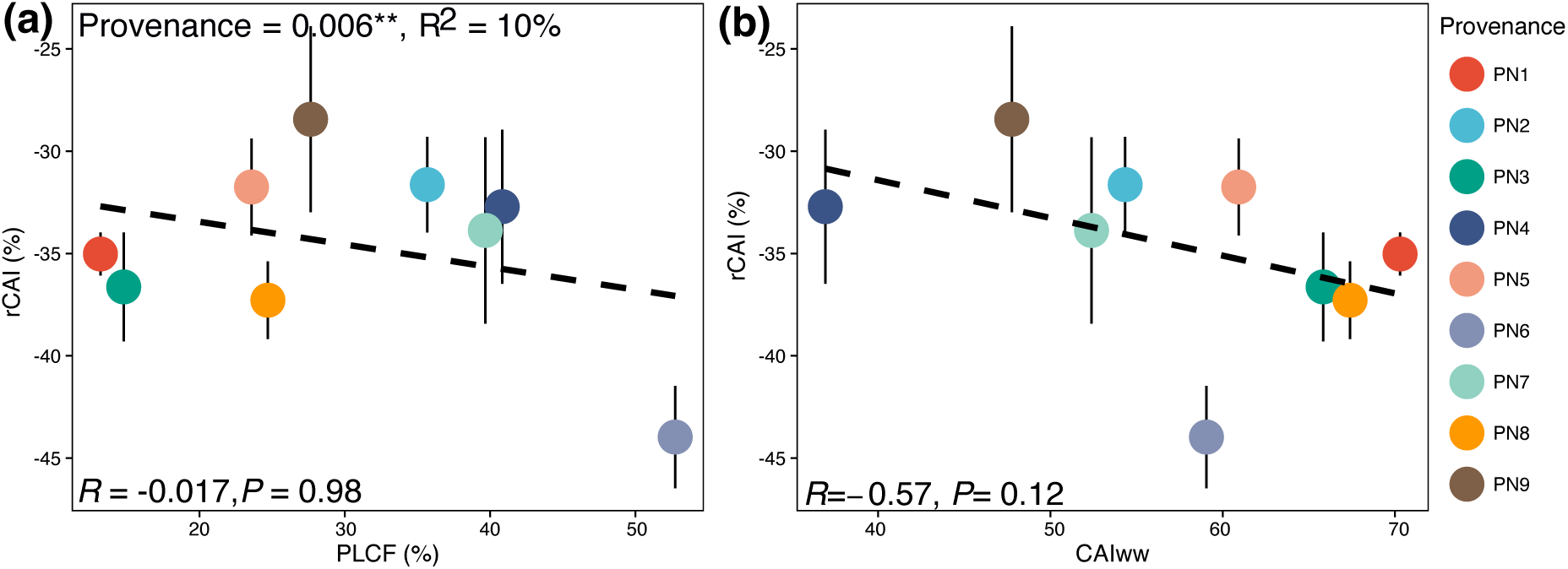
Effect of drought on relative canopy area increment (rCAI) and its relationship with drought tolerance across nine provenances of *Pinus nigra*. (a) rCAI plotted against drought tolerance measured as percentage loss of chlorophyll fluorescence (PLCF, %). (b) rCAI plotted against canopy area increment under well-watered conditions (CAI_WW_, %). Each point represents provenance means ± SE (9-15), with colors indicating provenances. Regression lines (dashed) are shown with corresponding correlation coefficients (R) and significance values (P). Provenance effects were significant for rCAI (P = 0.006, R² = 0.10).

**Fig. S3.**
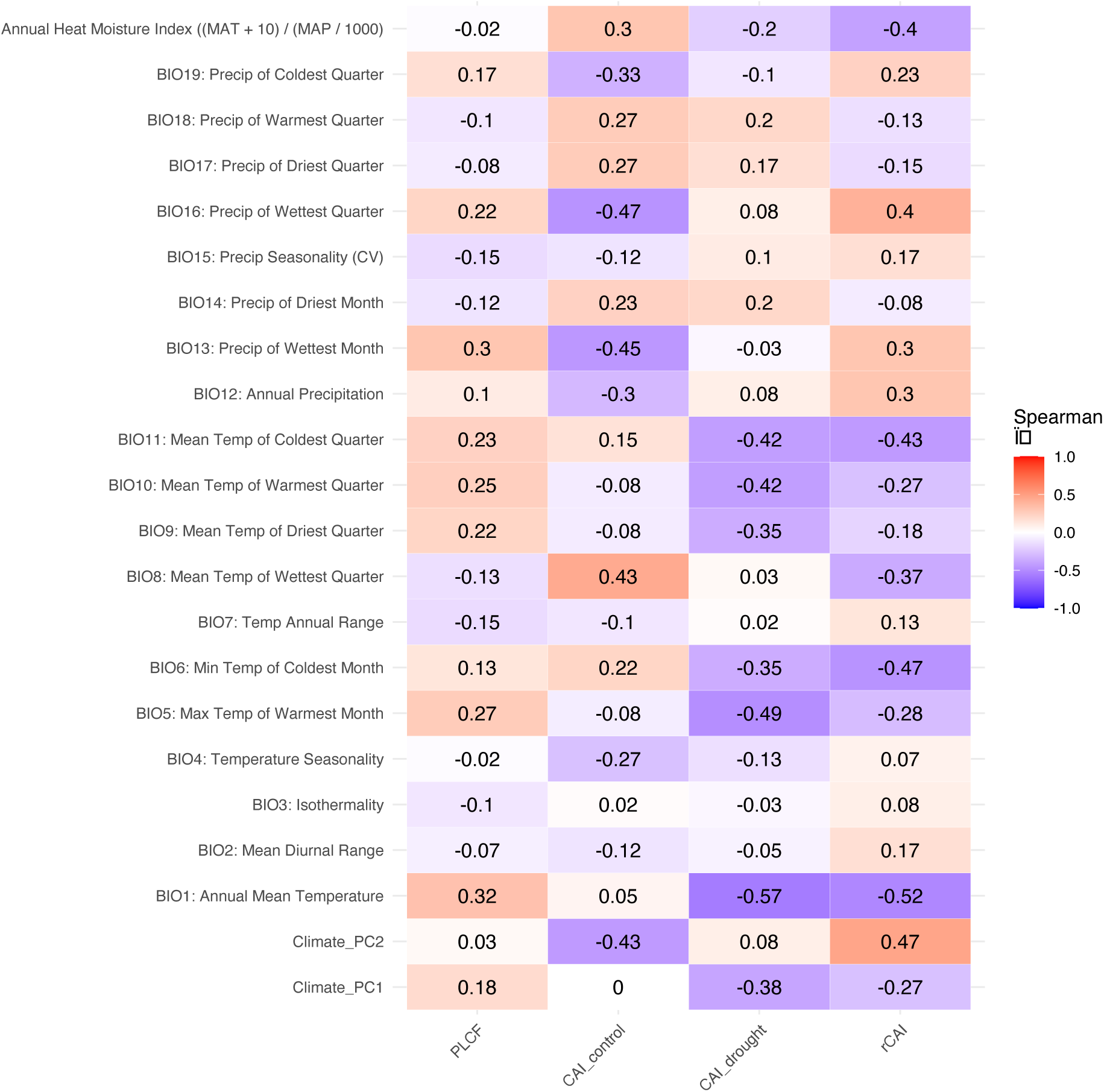
Spearman correlation between canopy area increment (CAI), relative-CAI (rCAI) or drought tolerance assessed based on percentage loss in chlorophyll fluorescence (PLCF) and 20 climatic variables from the site of origin of provenances. Positive and negative correlations (ρ) are represented by the color scale from blue to red, with numeric values indicating the Spearman correlation coefficient. Climate variables are sorted by grouping (temperature, precipitation, and derived indices).

**Fig. S4.**
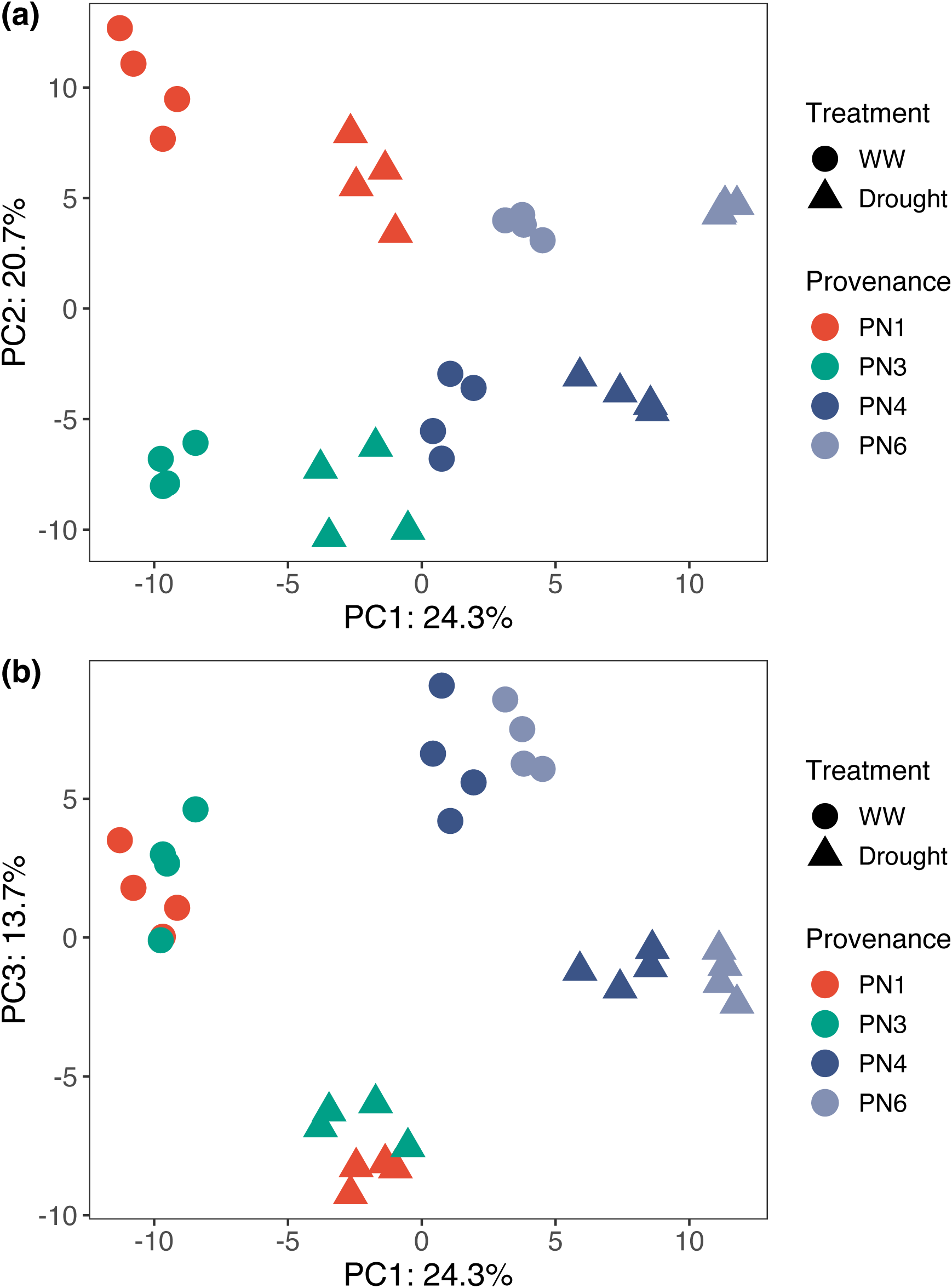
Principal component analysis (PCA) of metabolic profiles in *Pinus nigra* provenances under well-watered (WW) and drought conditions. (a) PC1 and PC2. (b) PC1 and PC3. PN1 and PN3 are drought tolerant (DT) while PN4 and PN6 are drought sensitive (DS) based on percentage loss in chlorophyll fluorescence (PLCF).

**Fig. S5.**
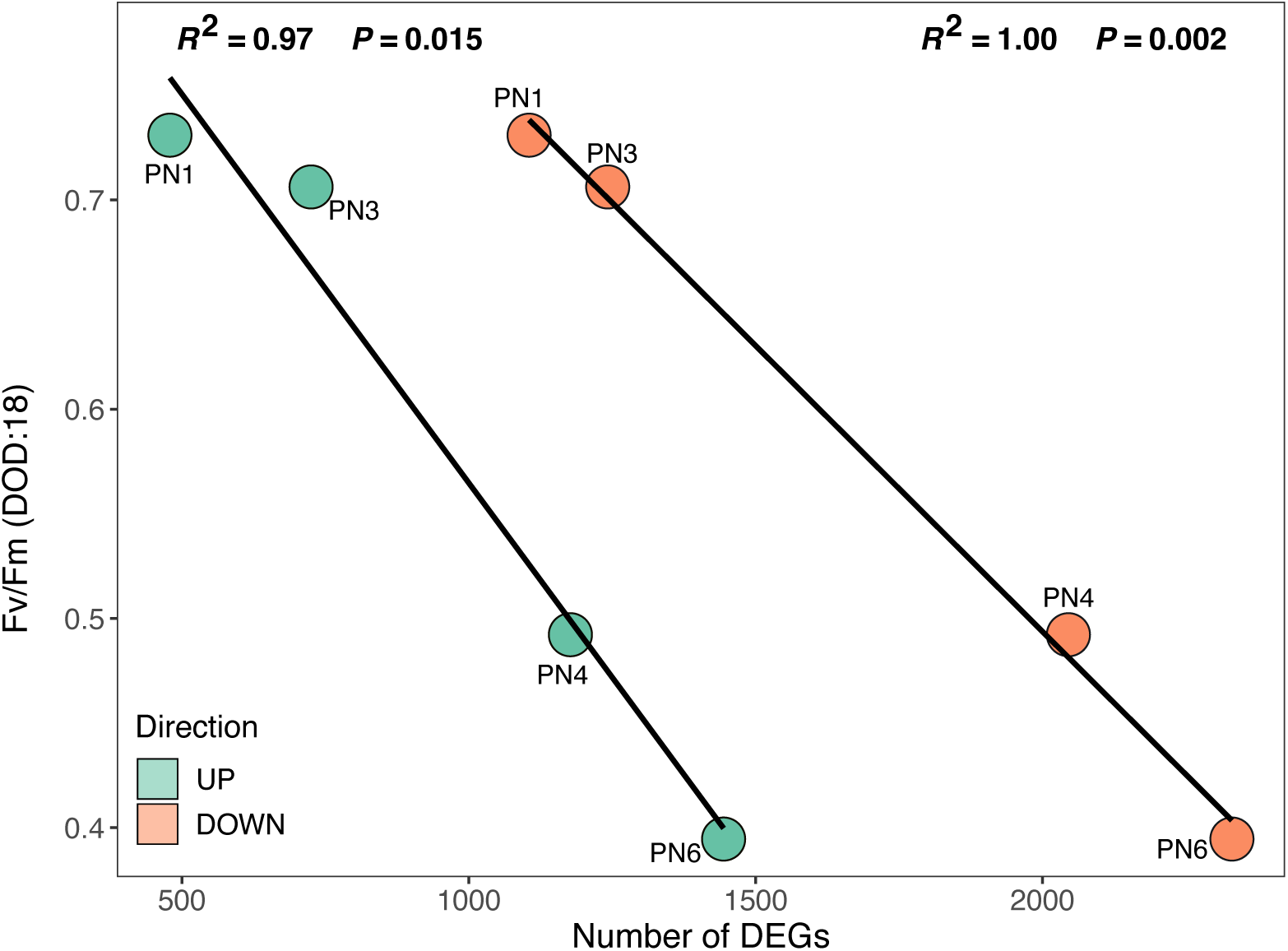
Relationship between the number of differentially expressed genes (DEGs) and maximum photochemical efficiency (Fv/Fm) on day 18 of drought. Each point represents a provenance, with colors indicating the direction of gene regulation (green for upregulated, orange for downregulated). Linear regressions are shown separately for up- and downregulated genes, with corresponding R² and P-values annotated.

**Fig. S6.**
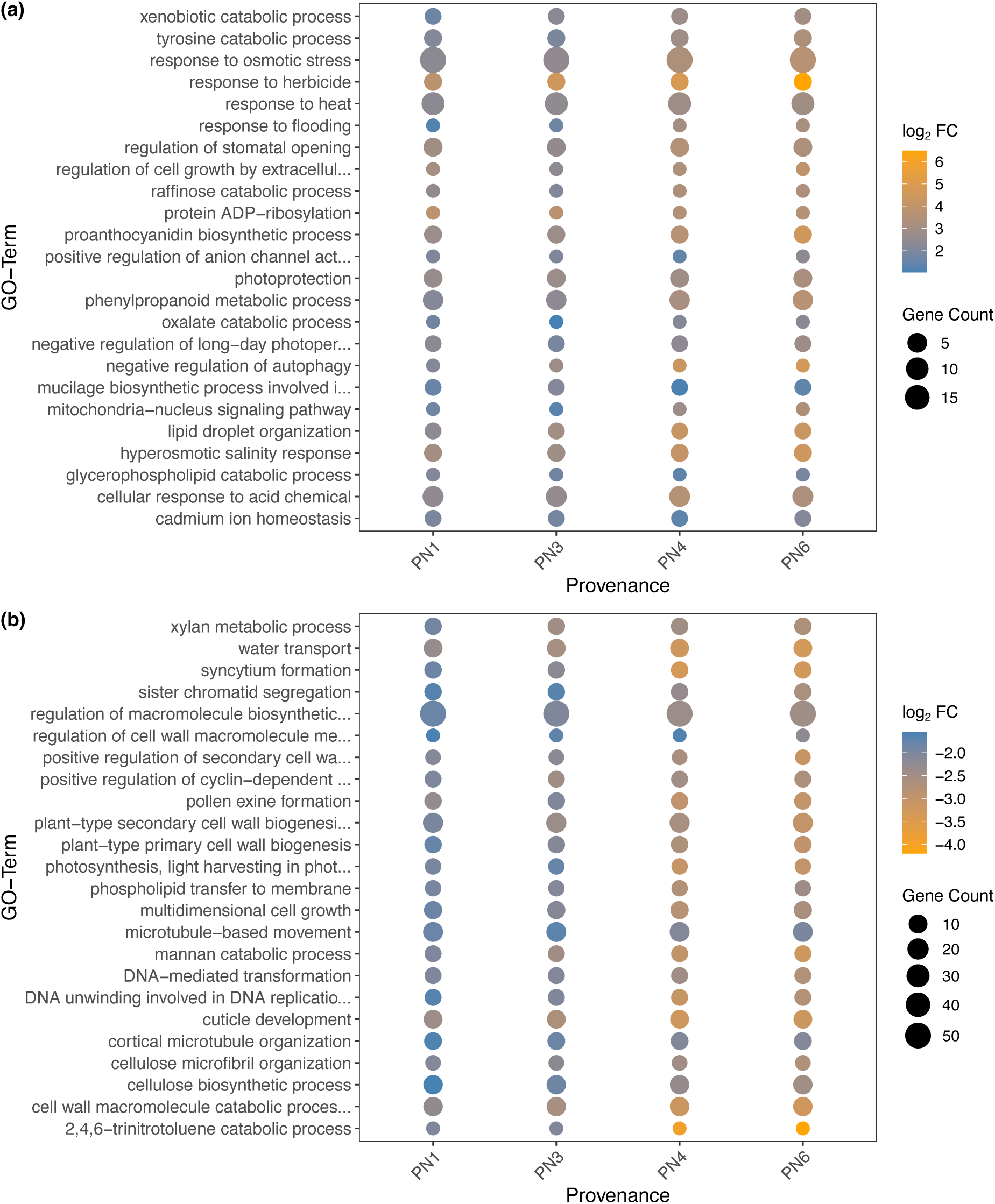
Gene Ontology (GO) enrichment analysis of differentially expressed genes (DEGs) across four *Pinus nigra* provenances under drought conditions. (a) GO terms enriched (P < 0.01) among upregulated genes. (b) GO terms enriched (P < 0.01) among downregulated genes. Each dot represents a GO term enriched in a given provenance. Dot size corresponds to the number of genes associated with the GO term (Gene Count), while color indicates the average log₂ fold change (log₂ FC) of genes in the term, with warmer colors representing stronger induction (panel a) or repression (panel b).

