## Supplementary material for "Unravelling the intraspecific variation in drought responses in seedlings of European black pine (*Pinus nigra* J.F. Arnold)": Fig. S

### Supplementary figures

### Fig. S1-S6

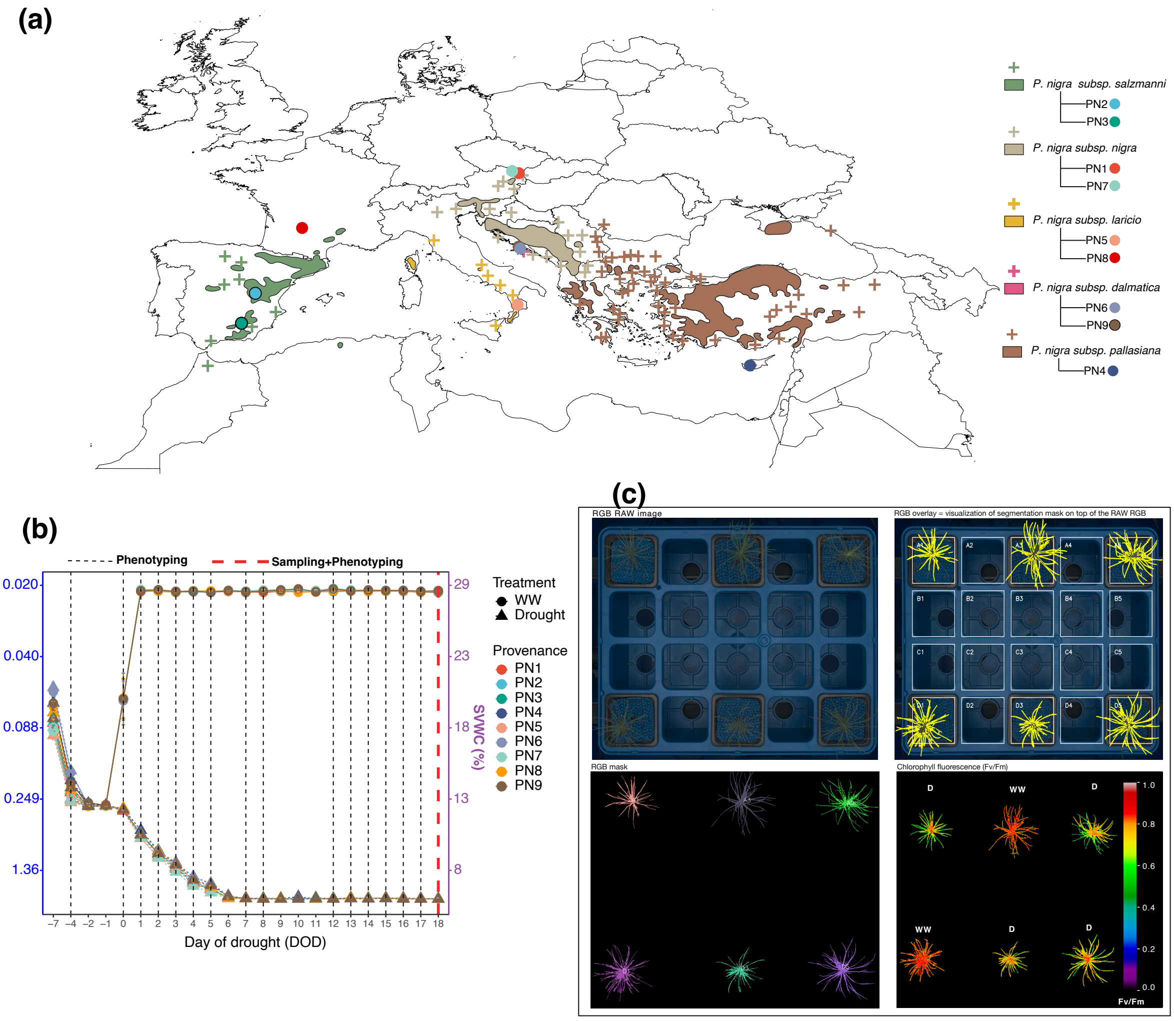


### Fig. S1

Study design. (a) Geographic origin of nine provenances representing five subspecies of black pine (*Pinus nigra*), shown alongside the natural distribution of each subspecies. (b) Drought progression measured as soil water potential over the 18-day drought stress experiment. Soil water potential decreased from -0.24 MPa to -2.97 MPa over six days and was maintained at this level for the remaining 12 days. (c) Schematic of the experimental set-up showing the imaging sensors used for high-throughput phenotyping. RGB: Red–Green–Blue imaging; Chlorophyll fluorescence imaging.

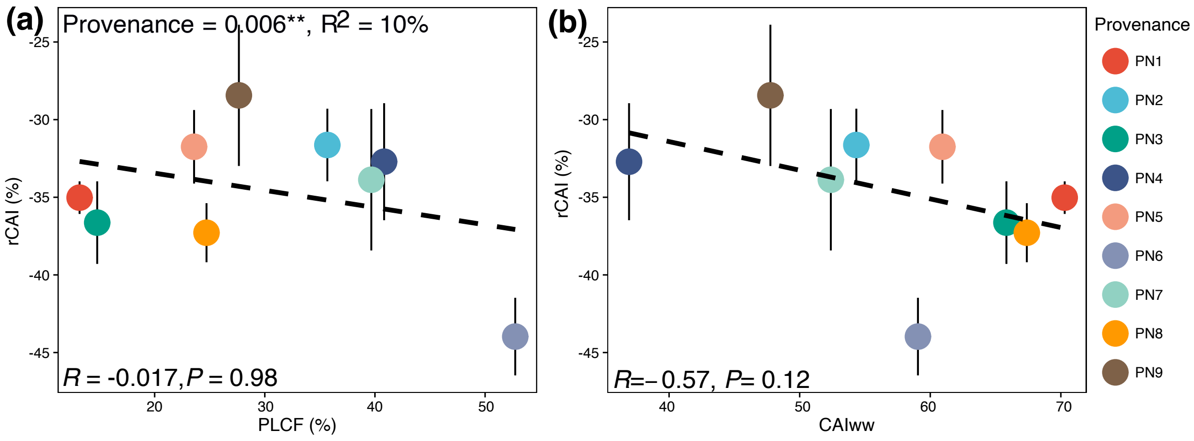


**Fig. S2**

Effect of drought on relative canopy area increment (rCAI) and its relationship with drought tolerance across nine provenances of *Pinus nigra*. (a) rCAI plotted against drought tolerance measured as percentage loss of chlorophyll fluorescence (PLCF, %). (b) rCAI plotted against canopy area increment under well-watered conditions ($CAI_{WW}$, %). Each point represents provenance means ± SE (9-15), with colors indicating provenances. Regression lines (dashed) are shown with corresponding correlation coefficients (R) and significance values (P). Provenance effects were significant for rCAI (P = 0.006, $R^2$ = 0.10).

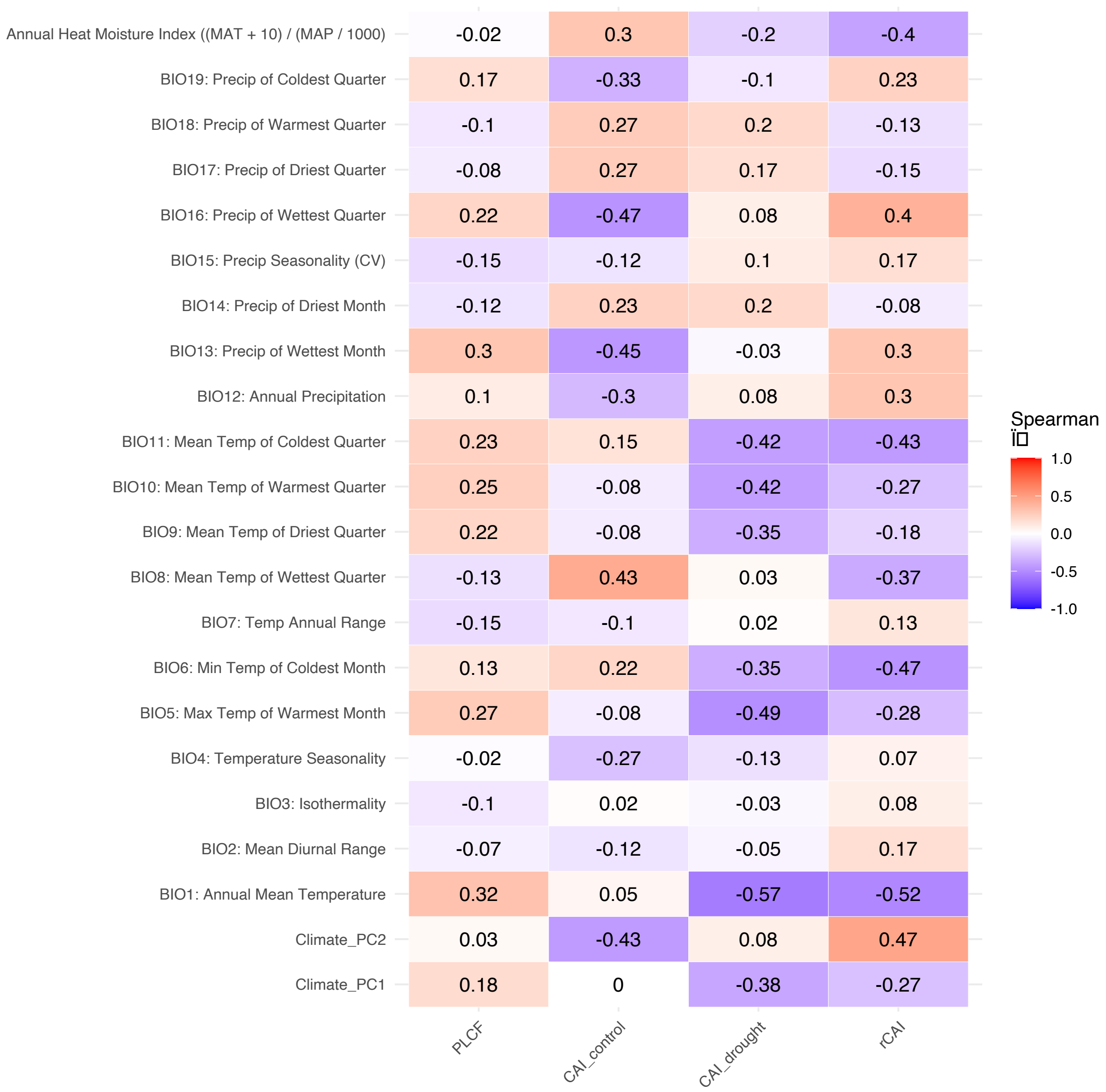


**Fig. S3**

Spearman correlation between canopy area increment (CAI), relative-CAI (rCAI) or drought tolerance assessed based on percentage loss in chlorophyll fluorescence (PLCF) and 20 climatic variables from the site of origin of provenances. Positive and negative correlations (ρ) are represented by the color scale from blue to red, with numeric values indicating the Spearman correlation coefficient. Climate variables are sorted by grouping (temperature, precipitation, and derived indices).

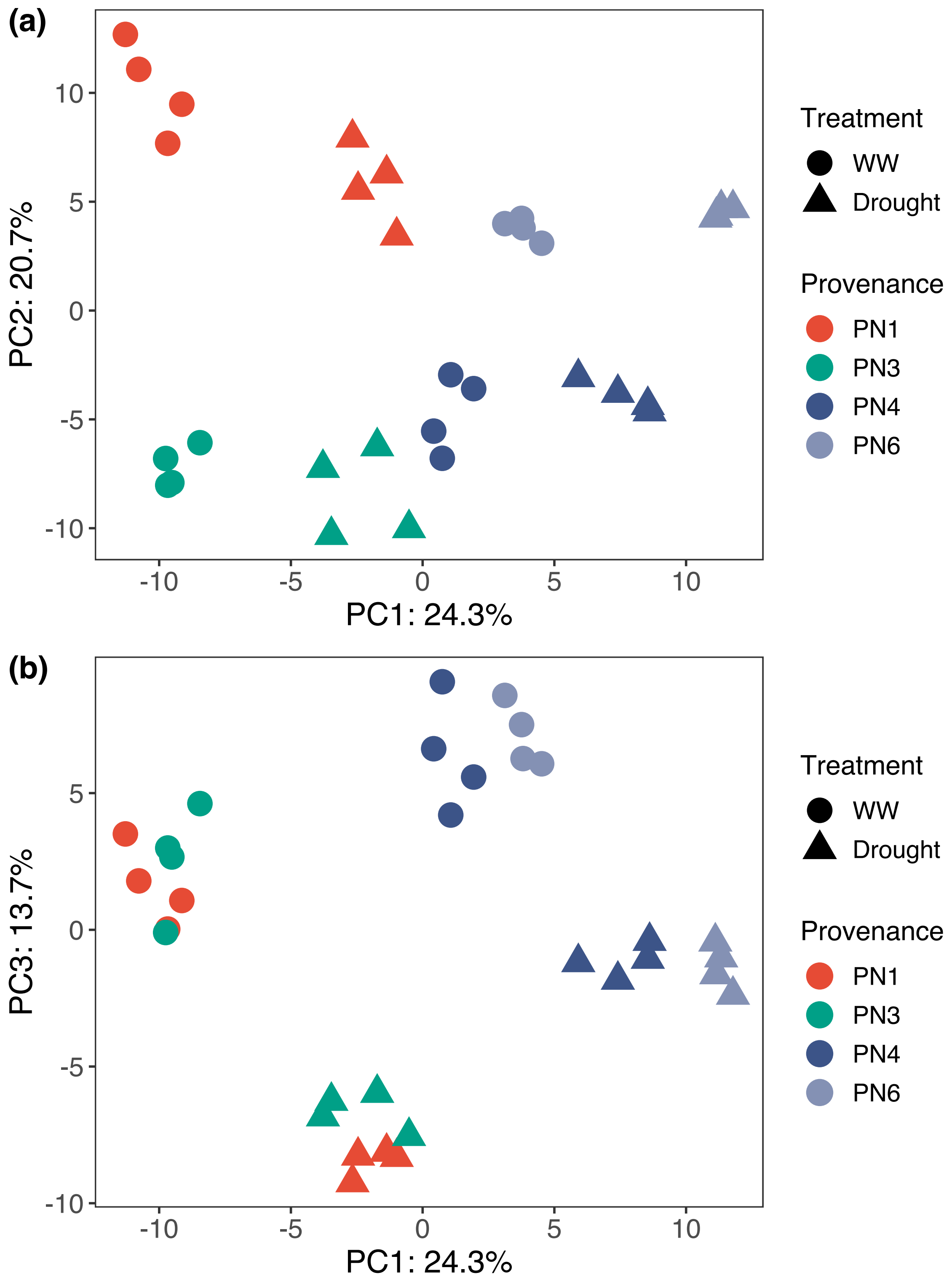


**Fig. S4**

Principal component analysis (PCA) of metabolic profiles in *Pinus nigra* provenances under well-watered (WW) and drought conditions. (a) PC1 and PC2. (b) PC1 and PC3. PN1 and PN3 are drought tolerant (DT) while PN4 and PN6 are drought sensitive (DS) based on percentage loss in chlorophyll fluorescence (PLCF).

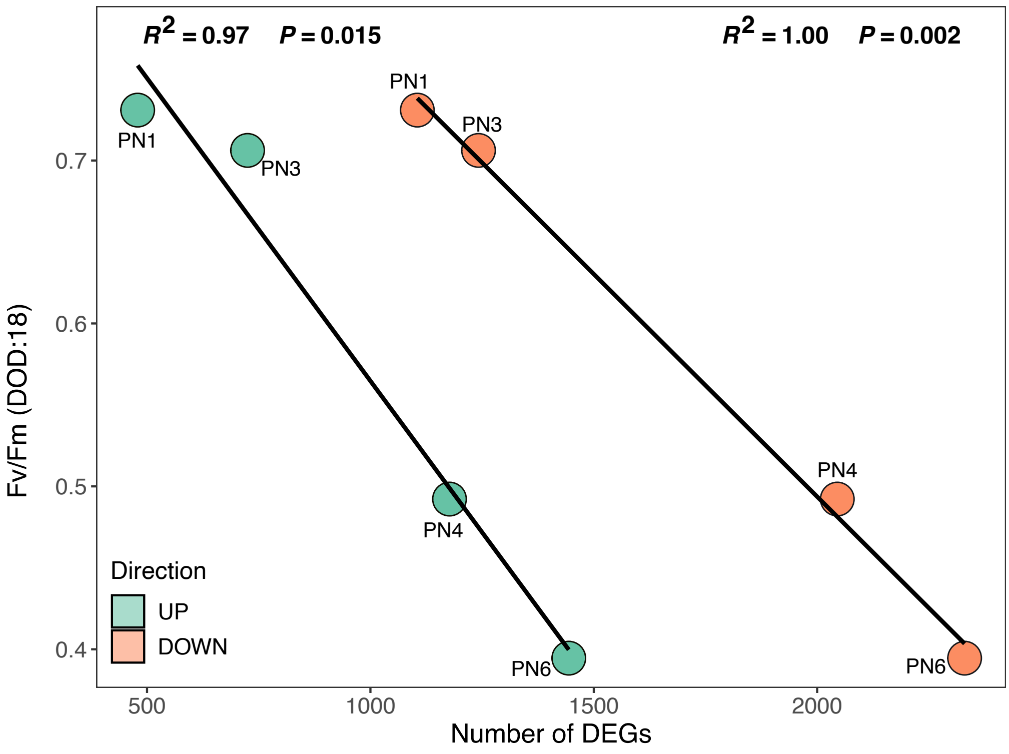


**Fig. S5**

Relationship between the number of differentially expressed genes (DEGs) and maximum photochemical efficiency (Fv/Fm) on day 18 of drought. Each point represents a provenance, with colors indicating the direction of gene regulation (green for upregulated, orange for downregulated). Linear regressions are shown separately for up- and downregulated genes, with corresponding $R^2$ and P-values annotated.

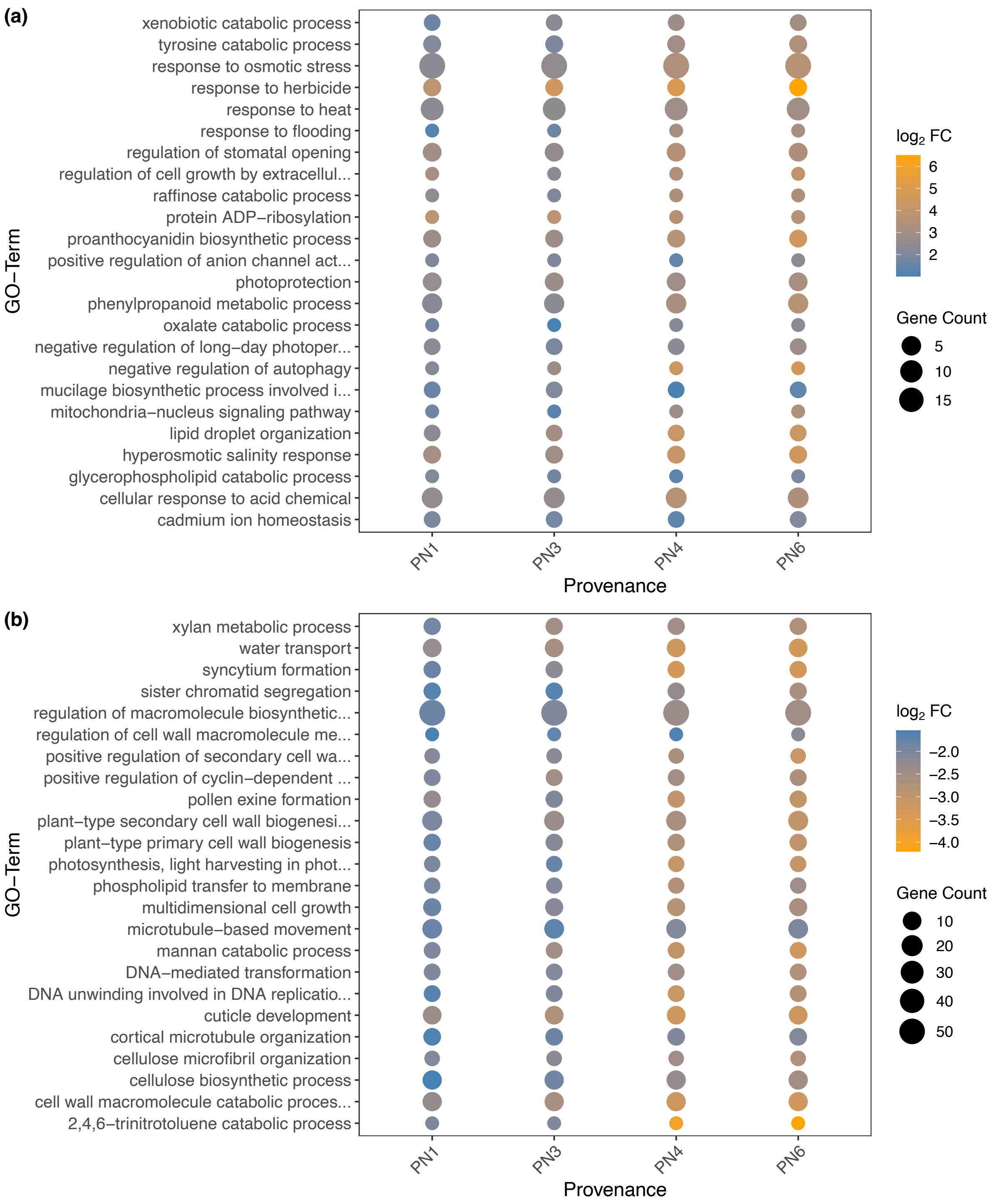


**Fig. S6**

Gene Ontology (GO) enrichment analysis of differentially expressed genes (DEGs) across four *Pinus nigra* provenances under drought conditions. (a) GO terms enriched ($P < 0.01$) among upregulated genes. (b) GO terms enriched ($P < 0.01$) among downregulated genes. Each dot represents a GO term enriched in a given provenance. Dot size corresponds to the number of genes associated with the GO term (Gene Count), while color indicates the average $\log_2$ fold change ($\log_2$ FC) of genes in the term, with warmer colors representing stronger induction (panel a) or repression (panel b).
