## Supplementary material for "Unravelling the intraspecific variation in drought responses in seedlings of European black pine (*Pinus nigra* J.F. Arnold)": Methods S1

**Supplementary materials**

**Methods S1**

***Image Analysis***

*RGB*

Images were read using the imread function within scikit-image version 0.19.1 (Van Der Walt *et al.*, 2014). Binary plant masks were created by transforming images to the L*a*b* color space using the cvtColor function within OpenCV version 4.5.5.62 and thresholding pixels in the ranges 0-255, 0-125, and 121-255 for the L*, a*, b* channels, respectively.

Objects with an area of less than 200 pixels were removed from the masks. Objects within the masks were identified using the findContours function in OpenCV. Contours were assigned to individual plants using the image coordinates of each tray location provided by the Plant Screen system. Plant area was calculated by summing the pixel area of all contours assigned to individual plants. Pixel areas were converted to mm^2^ using the pixel scale provided by the Plant Screen system.

*Chlorophyll fluorescence images*

Parameter image files were read as raw bites, converted to float values, and re-shaped into a 2D array corresponding to the image dimensions. Binary plant masks were created by thresholding pixels with a value greater than 700 in the maximum fluorescence (Fm) parameter images. Individual plants were assigned as described above. Mean parameter values for each plant were calculated by applying the binary mask to Fo and Fm image. Fv/Fm values were then estimated using formula (Fm-Fo)/Fm. For both the RGB and Chlf imaging, we removed any individual where images because of technical and quality issues could not be processed resulting in 14-15 and 9 individuals per provenance per treatment for PN1-PN8 and PN9, respectively.

**Extraction and quantification of soluble carbohydrates, condenses tannins and total phenolics**

For the measurement of metabolites from the same extract, we employed a sequential extraction protocol as described in (Preiner *et al.*, 2024). Briefly, all photometric assays were conducted using an extract from 15 mg of lyophilized and homogenized needle tissues per sample as described previously (Preiner *et al.*, 2024) with minor adaptions of standard concentrations and prior sample dilution to optimize for the concentration of metabolites. All chemicals were purchased from Sigma Aldrich, and absorbances of the assays were measured spectrophotometrically in triplicates in microplates (Greiner Bio-One, 96-Well microplate F-bottom). For quantification of soluble carbohydrates, a spectrophotometer of the type Multiskan® Spectrum (Thermo Scientific) was used, while for all other assays, we employed an Infinite® M Nano (Tecan) spectrophotometer. Extraction, as well as quantification of total phenolics, and condensed tannins, followed the procedures as described in (Preiner *et al.*, 2024). Samples were diluted prior to the assays for quantification of soluble carbohydrates (1:4). Concentrations of standard dilution curves were adjusted to cover the range of absorbances of samples in the assays for optimal calibration and absolute quantification of metabolite classes.

**Extraction and quantification of proline**

Proline content was determined in 25 mg aliquots of ground lyophilized material according to (Carillo *et al.*, 2008). Briefly, the samples were extracted twice at 4°C overnight with a total volume of 1.5 mL 40% EtOH. Proline was determined in triplicate 50 µL aliquots by ninhydrin staining and quantified based on standards of L-proline ranging from 0.04 to 1 mM.

**Extraction and quantification of flavonoids**

Lyophilized finely ground plant material (25 mg) was extracted for 2 h with 1.6 mL 80% analytical grade methanol with 0.1 mg/mL 2-phenylethanol as an internal standard. The extracts were centrifuged at maximum speed. High performance liquid chromatography (HPLC) analyses were conducted using an Agilent 1200 instrument equipped with a Bruker Daltronics (Germany) 2010 model photodiode array detector. Twenty μl extract was injected on the chromatographic column (EC 250 × 4.6 mm NUCLEODUR Sphinx RP, 5 μm, Macherey Nagel, Germany) equipped with a precolumn (C18, 5 μm, 4 × 3 mm, Phenomenex, USA). The mobile phase consisted of HPLC-grade water amended with 0.2% formic acid (FA) and acetonitrile (ACN). The flow rate was set to 0.8 mL/min and the column was maintained at 25 °C. The gradient was as follows: 0 min, 95% water/FA; 0 to 18 min, 5% to 65% ACN; 18 to 22 min, 95% ACN; 22 to 26 min, 95% water/FA. Flavonoids were detected by absorption at 330 nm and flavan-3-ols at 280 nm. Compounds were identified by comparison of retention times in relation to those of standards, where available. Quantities were calculated on the basis of peak areas using a taxifolin standard curve corrected by the recovery of the internal standard and experimentally determined response factors (3.7 for flavan-3-ols, 1.1 for quercetin and 1.3 for kaempferol). Putative identities for flavonoid glycosides were assigned based on ion masses obtained on a Bruker Datronics Esquire 3000 ion trap operated in negative mode following the methods described by (Hammerbacher *et al.*, 2018).
